# Disease-associated astrocyte epigenetic memory promotes CNS pathology

**DOI:** 10.1101/2024.01.04.574196

**Authors:** Hong-Gyun Lee, Joseph M. Rone, Zhaorong Li, Camilo Faust Akl, Seung Won Shin, Joon-Hyuk Lee, Lucas E. Flausino, Florian Pernin, Chun-Cheih Chao, Kilian L. Kleemann, Lena Srun, Tomer Illouz, Federico Giovannoni, Marc Charabati, Liliana M. Sanmarco, Jessica E. Kenison, Gavin Piester, Stephanie E. J. Zandee, Jack Antel, Veit Rothhammer, Michael A. Wheeler, Alexandre Prat, Iain C. Clark, Francisco J. Quintana

## Abstract

Astrocytes play important roles in the central nervous system (CNS) physiology and pathology. Indeed, astrocyte subsets defined by specific transcriptional activation states contribute to the pathology of neurologic diseases, including multiple sclerosis (MS) and its pre-clinical model experimental autoimmune encephalomyelitis (EAE)^1–8^. However, little is known about the stability of these disease-associated astrocyte subsets, their regulation, and whether they integrate past stimulation events to respond to subsequent challenges. Here, we describe the identification of an epigenetically controlled memory astrocyte subset which exhibits exacerbated pro-inflammatory responses upon re-challenge. Specifically, using a combination of single-cell RNA sequencing (scRNA-seq), assay for transposase-accessible chromatin with sequencing (ATAC–seq), chromatin immunoprecipitation with sequencing (ChIP–seq), focused interrogation of cells by nucleic acid detection and sequencing (FIND-seq), and cell-specific *in vivo* CRISPR/Cas9-based genetic perturbation studies we established that astrocyte memory is controlled by the metabolic enzyme ATP citrate lyase (ACLY), which produces acetyl coenzyme A (acetyl-CoA) used by the histone acetyltransferase p300 to control chromatin accessibility. ACLY^+^p300^+^ memory astrocytes are increased in acute and chronic EAE models; the genetic targeting of ACLY^+^ p300^+^ astrocytes using CRISPR/Cas9 ameliorated EAE. We also detected responses consistent with a pro-inflammatory memory phenotype in human astrocytes *in vitro*; scRNA-seq and immunohistochemistry studies detected increased ACLY^+^ p300^+^ astrocytes in chronic MS lesions. In summary, these studies define an epigenetically controlled memory astrocyte subset that promotes CNS pathology in EAE and, potentially, MS. These findings may guide novel therapeutic approaches for MS and other neurologic diseases.

---

Astrocytes are abundant non-hematopoietic glial cells of the central nervous system (CNS) with important functions in health and disease^1–3^. Astrocytes participate in key processes relevant to CNS development and homeostasis^1^. In addition, cytokines, interactions with CNS-resident and CNS-recruited immune cells, and other factors trigger astrocyte responses with important roles in CNS pathology^2,4,5^. Indeed, several astrocyte subsets have been described in neurologic diseases^6–8^. For example, we and others interrogated astrocyte functional heterogeneity in multiple sclerosis (MS) and its model experimental autoimmune encephalomyelitis (EAE)^9–16^. However, the stability of these disease-associated astrocyte subsets is unclear, an important point when considering life-long chronic neurologic diseases such as MS.

Immunological memory, the generation of faster and stronger responses upon repeated antigenic stimulation, is a classic hallmark of adaptive immunity driven by long-lived antigen-specific T cells and B cells^17^. However, upon stimulation, innate immune cells including myeloid cells^18,19^, natural killer (NK) cells^20^, and innate lymphoid cells (ILCs)^21^ undergo metabolic, epigenetic, and transcriptional adaptations that alter their subsequent response to stimulation, boosting protective immunity against pathogens but also contributing to the pathogenesis of inflammatory diseases^22^. Although memory T cells and B cells have been identified, our understanding of innate immune or non-hematopoietic cell memory subsets remains limited. In this context, it is still unknown whether astrocytes display altered responses to repeated stimulation, how these responses are regulated, and whether specific astrocyte subsets are involved.

Here, we describe a memory astrocyte subset controlled by epigenetic changes driven by ACLY- and p300-dependent histone acetylation, which following an initial stimulation display faster and stronger pro-inflammatory responses upon restimulation. This subset of memory astrocytes is expanded in EAE and MS; its genetic inactivation ameliorates EAE. These findings highlight important mechanisms relevant to the pathology of MS as well as other chronic neurologic disorders and may guide the development of novel therapeutic interventions.

## Astrocyte epigenetic memory

To investigate whether an initial pro-inflammatory stimulus affects subsequent astrocyte responses, we administered IL-1β and TNF (intracerebroventricular, ICV) to mice once (1X IL-1β+TNF), or twice one week apart (2X IL-1β+TNF); vehicle was used as a control (**Fig. 1a**). These cytokines were selected because they trigger astrocyte responses similar to those detected in MS and EAE^15^. One day after the last stimulation, we analyzed by RNA-seq brain astrocytes from the 1X and 2X treatment groups. Based on the number and expression levels of differentially expressed genes (DEGs), 2X IL-1β+TNF stimulation induced more potent pro-inflammatory astrocyte responses than 1X IL-1β+TNF (**Figs. 1b**-**d**, **Extended Data Figs. 1a,b, Supp. Table 1)**.

**Figure 1:**
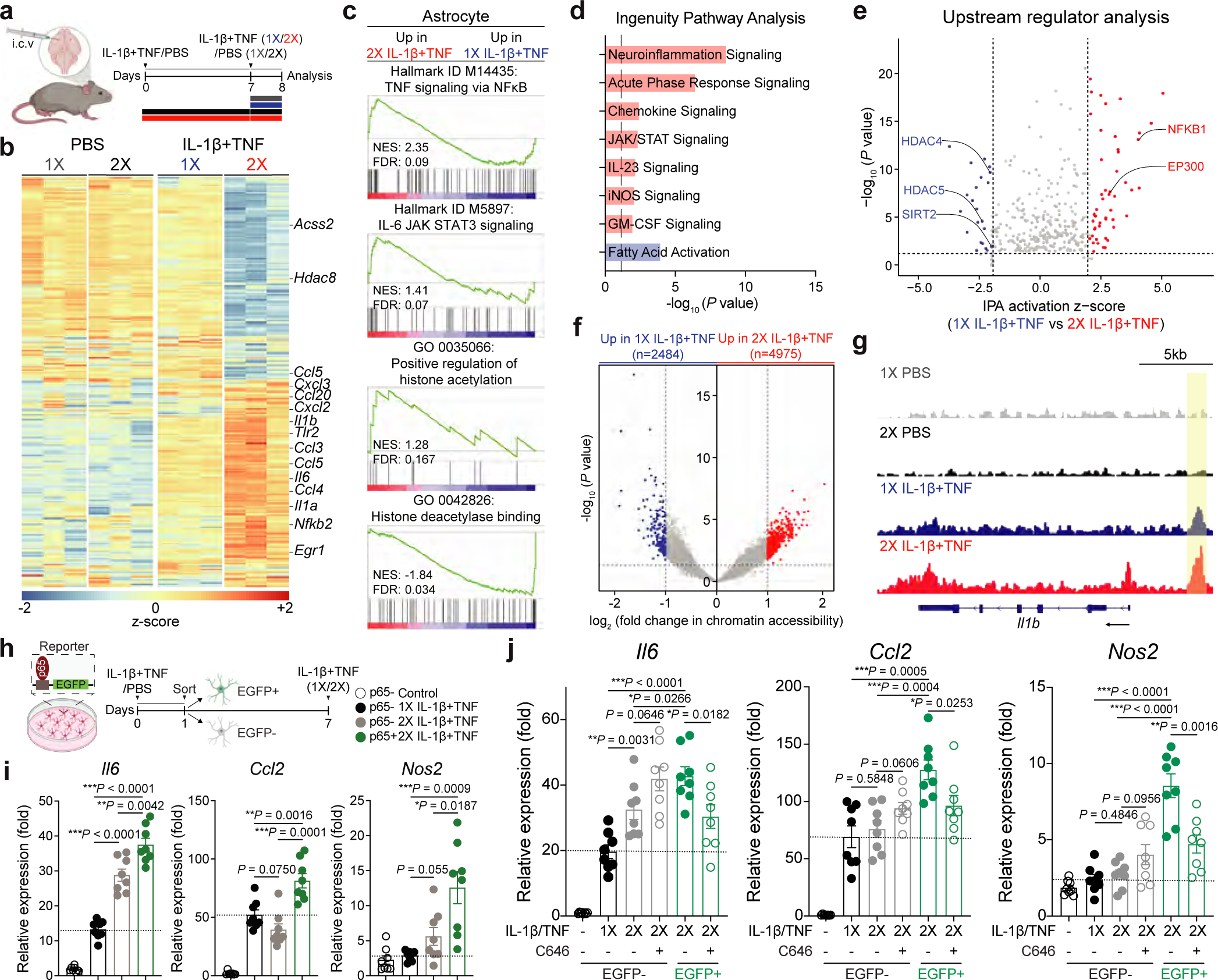
Pro-inflammatory stimuli induce astrocyte epigenetic memory. **(a)** Experimental design for (b) to (g). C57BL/6 mice received ICV administration of IL-1β/TNF or PBS once (1X) or twice (2X). After 18h, sorted brain astrocytes were analyzed. **(b)** Differential gene expression determined by RNA-seq in sorted astrocytes (n=3 per group). **(c)** Gene set enrichment analysis (GSEA) comparing astrocytes stimulated twice (2X) versus once (1X). Gene Ontology (GO) terms are shown. FDR, false discovery rate; NES, normalized enrichment score. **(d,e)** IPA (left) and predicted upstream regulators (right) in isolated astrocytes. Up-(red) or down-regulated (blue) in astrocytes stimulated twice (2X) versus once (1X) are shown. **(f,g)** ATAC-seq analysis of astrocytes comparing 2X IL-1β+TNF versus 1X IL-1β+TNF (n=5 per group). **(f)** Genes depicting increased (red) or decreased (blue) accessibility in astrocytes. **(g)** Genome browser snapshots showing the ATAC-seq sequencing tracks at the *Il1b* locus. Only regions showing a significant increase (*p*-value < 0.05) in accessibility in astrocytes from mice stimulated twice (2X) versus once (1X) are highlighted by yellow boxes. **(h)** Experimental design for (i). Primary astrocytes isolated from *p65^EGFP^* reporter mice received PBS or IL-1β/TNF stimulation. After 18-24h, EGFP positive/negative astrocytes were sorted and received IL-1β/TNF stimulation once (1X) or twice (2X) after 6 days. **(i)** qPCR of EGFP positive/negative astrocytes after 30min stimulation with IL-1β/TNF on day 7 (n=8 per group). Unpaired two-sided *t*-test. **(j)** Gene expression 30min after IL-1β/TNF stimulation on day 7 of EGFP positive/negative astrocytes treated with C646 or vehicle (n=8 per group). Unpaired two-sided *t*-test. Data shown as mean ± s.e.m.

Ingenuity Pathway Analysis (IPA) identified NF-κB as an upstream regulator of the astrocyte response to IL-1β+TNF. However, NF-κB signaling was higher in the 2X IL-1β+TNF group (**Fig. 1e**). In addition, 2X IL-1β+TNF treatment also increased transcriptional responses linked to histone acetylation, while decreasing transcriptional responses linked to histone deacetylase (HDAC) activity (**Fig. 1c**). Indeed, we also identified *Ep300*, which encodes histone acetyltransferase (HAT) p300, as an upstream regulator of the transcriptional response of astrocytes to 2X IL-1β+TNF stimulation (**Fig. 1e**, **Extended Data Fig. 1c**). In support of these findings, the assay for transposase-accessible chromatin with sequencing (ATAC)-seq detected increased chromatin accessibility in inflammation- and epigenetic regulation-related genes in the 2X IL-1β+TNF treatment group (**Figs. 1f**,**g**, **Extended Data Figs. 1d,e, Supp. Table 2)**. These genes depicting increased chromatin accessibility displayed motif enrichment for NF-κB, ATF-1, and ATF-3, transcription factors known to interact with p300 and its paralog CREB-binding protein (CBP)^23,24^ **(Extended Data Fig. 1f)**.

These *in vivo* observations may reflect changes in astrocyte-intrinsic responses and/or the control of astrocytes by other cells such as microglia, known to modulate astrocyte responses^2,5^. To focus on astrocyte-intrinsic mechanisms, we used primary mouse brain astrocyte cultures. Astrocyte stimulation two times one week apart (2X IL-1β+TNF) boosted genes associated with astrocyte proinflammatory activities (*Il6*, *Ccl2*, and *Nos2*)^10,16^ and decreased neuronal viability when compared to 1X IL-1β+TNF stimulation **(Extended Data Figs. 1g-i)**. The 2X IL-1β+TNF treatment group also depicted the largest decrease in lactate release **(Extended Data Fig. 1j)**, suggestive of a stronger impairment in astrocyte capacity to support neuron metabolic needs^25^. Of note, astrocyte pro-inflammatory gene expression induced by IL-1β+TNF stimulation *in vitro* returned to basal levels 7 days after the initial cytokine stimulation (**Extended Data Fig. 1k)**.

The increased responsiveness detected following 2X IL-1β+TNF stimulation of astrocytes in culture may reflect increased cell-intrinsic responses in previously activated astrocytes and/or the engagement of additional astrocytes not activated in the first activation round. To investigate the contribution of cell-intrinsic responses in previously activated astrocytes, we used primary mouse brain astrocyte cultures derived from *p65^EGFP^* reporter mice. In these astrocyte cultures, IL-1β+TNF stimulation induced p65 nuclear translocation and EGFP expression **(Extended Data Fig. 1l)**; 2X IL-1β+TNF stimulation triggered more NF-kB activation than 1X IL-1β+TNF. Thus, we isolated EGFP^+^ and EGFP^-^ astrocytes by FACS following one round of IL-1β+TNF stimulation, and after a resting period of 7 days, we restimulated EGFP^+^ and EGFP^-^ sorted astrocytes with IL-1β+TNF (**Fig. 1h**, **Extended Data Fig. 1m)**. Please note that both EGFP and pro-inflammatory gene expression returned to basal levels 7 days after the first stimulation with IL-1β+TNF **(Extended Data Figs. 1n,o)**. We detected higher *Il6*, *Ccl2,* and *Nos2* expression upon restimulation, as well as stronger NF-kB activation, in EGFP^+^ versus EGFP^-^ sorted astrocytes (**Fig. 1i**, **Extended Data Fig. 1p)**. These findings suggest that cell-intrinsic mechanisms alter the subsequent response of a subset of astrocytes to stimulation.

Our RNA-seq and ATAC-seq analyses detected increased transcriptional signatures linked to histone acetylation in the 2X IL-1β+TNF treatment group (**Fig. 1c**, **Extended Data Figs. 1c,f)**, identifying HATs p300, Tip60, and PCAF as candidate regulators **(Extended Data Figs. 1c, 2a)**. However, in primary mouse astrocytes activated *in vitro* with 2X IL-1β+TNF, we detected an increase in *Ep300*, but not *Kat5* (Tip60) or *Kat2b* (PCAF) expression; increased *Ep300* expression persisted upon additional stimulation with IL-1β+TNF (3X) **(Extended Data Figs. 2b,c)**. Indeed, the p300 inhibitor C646 suppressed the increase of *Il6*, *Ccl2*, and *Nos2* expression induced by 2X IL-1β+TNF **(Extended Data Figs. 2d,e)**. Of note, pharmacological inactivation of p300 did not alter NF-kB expression or activation following 1X IL-1β+TNF stimulation, but abrogated the increased expression of *Il6, Ccl2*, and *Nos2* detected after 2X IL-1β+TNF stimulation in sorted EGFP^+^ *p65^EGFP^*reporter astrocytes (**Fig. 1j**, **Extended Data Figs. 2f,g)**. Taken together, these findings suggest that p300-driven epigenetic memory boosts astrocyte pro-inflammatory responses following an initial pro-inflammatory stimulation.

## p300 drives astrocyte epigenetic memory

Astrocytes play important roles in the pathology of MS and its preclinical model EAE^9–16^. To study the relevance of astrocyte epigenetic memory *in vivo*, we first generated a transcriptional signature score based on the response to 2X IL-1β+TNF treatment detected by RNA-seq (**Fig. 1a**, **Extended Data Fig. 3a, Supp. Table 3)**. The analysis of our astrocyte scRNA-seq datasets^10^ detected an increased transcriptional signature score of epigenetic memory-driven astrocytes at the peak of EAE **(Extended Data Fig. 3b)**. Indeed, ICV IL-1β+TNF stimulation of brain astrocytes isolated 22 days after EAE induction induced higher *Il6*, *Ccl2,* and *Nos2* expression than in naïve mice, resembling the increased responses detected *in vitro* following 2X IL-1β+TNF **(Extended Data Figs. 2c-e)**.

Histone 3 acetylation at lysine 27 (H3K27ac) is controlled by p300, which our studies identified as a candidate regulator of astrocyte epigenetic memory (**Fig. 1j**, **Extended Figs. 1c,f, 2e**). Indeed, we detected an increase in p300^+^ and H3K27ac^+^p300^+^ astrocytes in spinal cord white matter (WM) during EAE (**Fig. 2a**). To study the functional role of p300 expressed in astrocytes during EAE, we inactivated *Ep300* using a *Gfap* promoter-driven CRISPR/Cas9 lentiviral vector as previously described^9–12,14,15^. *Ep300* inactivation in CNS astrocytes ameliorated EAE, reduced demyelination, and suppressed pro-inflammatory astrocyte responses as determined by RNA-seq but did not affect CNS-recruited or peripheral CD4^+^ T cells (**Figs. 2b**-**d**, **Extended Data Figs. 3f-k, Supp. Table 4**). *Ep300* inactivation also suppressed the transcriptional signature linked to epigenetic memory astrocytes (**Figs. 2c,d**), and reduced NF-κB signaling and H3K27ac expression compared to scramble control **(Extended Data Figs. 3f,h)**. In support of these findings, chromatin immunoprecipitation followed by sequencing (ChIP-seq) detected decreased H3K27ac marks in genes related to metabolism, NF-κB signaling, and inflammation following *Ep300* inactivation in astrocytes (**Figs. 2e**-**g**, **Extended Fig. 3l, Supp. Table 5)**. Further support for a role of p300 in the control of astrocyte epigenetic memory was provided by the analysis of primary mouse brain astrocytes in culture, where 2X IL-1β+TNF boosted p300 recruitment to *Il6, Ccl2,* and *Nos2* promoters **(Extended Data Fig. 3m)**. Collectively, these findings suggest that p300-driven histone acetylation regulates pro-inflammatory epigenetic memory astrocyte responses in EAE.

**Figure 2:**
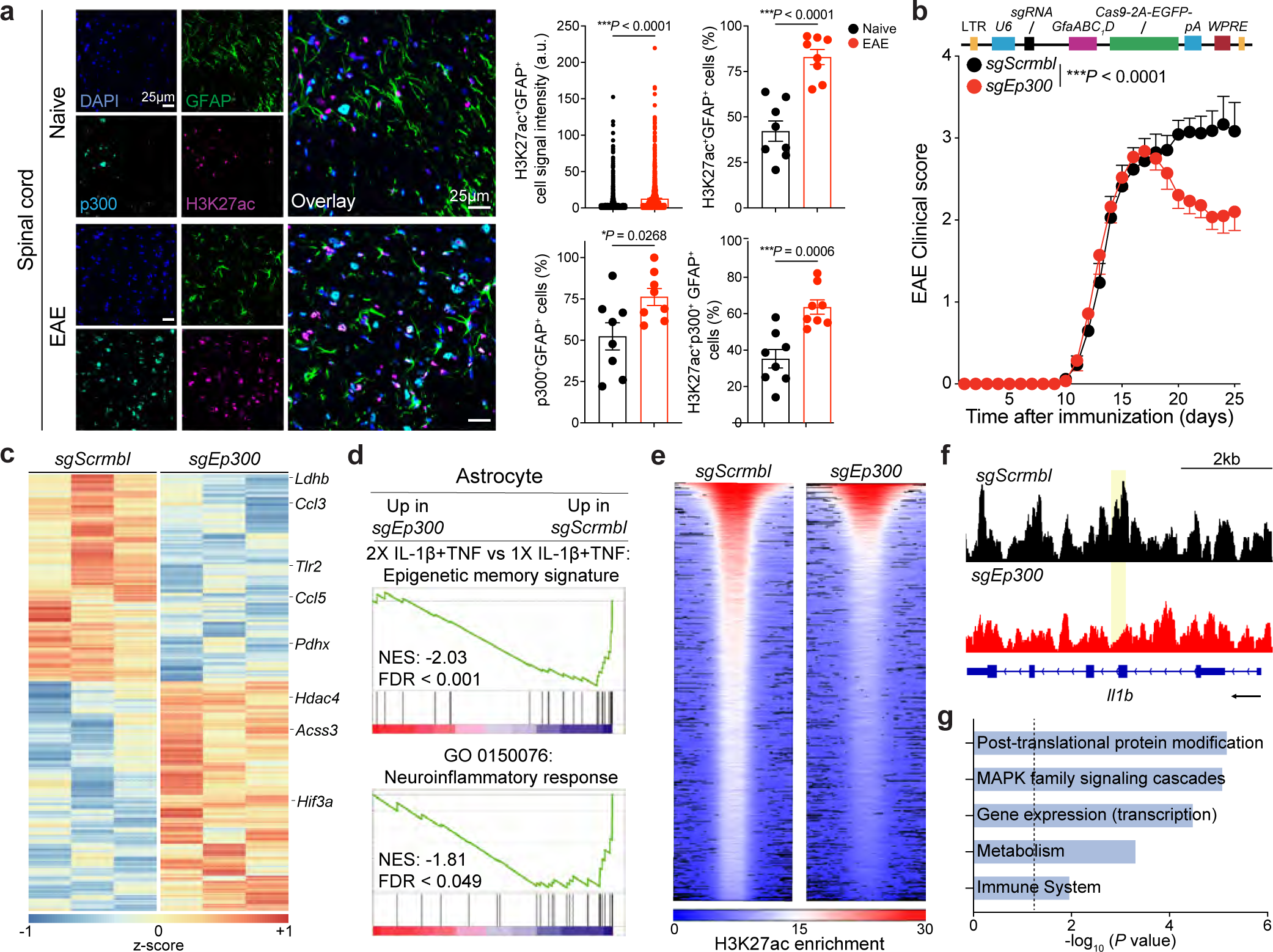
p300 promotes astrocyte epigenetic memory in EAE. **(a)** Immunostaining (left) and quantification (right) of H3K27ac^+^ and p300^+^ astrocytes in mice with or without EAE (n=8 spinal cord sections; n=3 mice per group). Astrocyte H3K27ac levels were calculated as the mean signal intensity (arbitrary units) per GFAP+ cells using automated unbiased quantification. Unpaired two-sided *t*-test. **(b)** EAE curves in sg*Scrmbl* (n=17) and sg*Ep300* (n=14) mice. Representative data of three independent experiments. Two-way repeated measures of ANOVA. **(c)** Differential gene expression determined by RNA-seq in astrocytes from sg*Scrmbl*- and sg*Ep300*-transduced mice 23 days after EAE induction (n=3 per group). **(d)** GSEA comparing sg*Scrmbl*- and sg*Ep300*-transduced astrocytes. **(e-f)** Chip-seq analysis of sg*Scrmbl*- and sg*Ep300*-transduced astrocytes (n=3 per group). **(e)** Heat map showing dynamic H3K27 acetylation marks. **(f)** Genome browser snapshots showing the *Il1b* locus. Only regions showing a significant decrease (*p*-value < 0.05) in accessibility in sg*Scrmbl*-transduced versus sg*Ep300*-transduced astrocytes are highlighted by yellow boxes. **(g)** Reactome pathway analysis down-regulated (blue) in Chip-seq accessible peaks of isolated astrocytes comparing sg*Scrmbl*-transduced versus sg*Ep300*-transduced astrocytes. Data shown as mean ± s.e.m.

## ACLY controls p300-driven astrocyte epigenetic memory

HATs acetylate lysine residues on histones using the substrate acetyl-coenzyme A (acetyl-CoA), a metabolite that controls metabolic and epigenetic networks^26^. In mammalian cells, the acetyl-CoA used for histone acetylation is generated by two major enzymes: ATP-citrate lyase (ACLY) and acetyl-CoA synthetase 2 (ACSS2), which use citrate and acetate as substrates, respectively^26,27^. To determine which enzyme provides the acetyl-CoA involved in the control of astrocyte epigenetic memory responses, we first analyzed primary mouse brain astrocytes stimulated *in vitro* with 1X or 2X IL-1β+TNF, finding that astrocyte re-stimulation boosts *Acly* but not *Acss2* expression **(Extended Data Fig. 4a)**. Indeed, *Acly* expression persisted upon additional IL-1β+TNF stimulation (3X), in alignment with our findings on *Ep300 expression* **(Extended Data Figs. 2c,4b)**. Moreover, in immunohistochemistry (IHC) studies, we detected an increase in the number of ACLY^+^ astrocytes in spinal cord WM during EAE, concomitant with a decrease in ACSS2^+^ astrocytes **(Extended Data Figs. 4c,d)**, suggesting a role for ACLY in the control of astrocyte epigenetic memory.

To evaluate the functional relevance of these findings, we inactivated *Acly* or *Acss2* in CNS astrocytes using a *Gfap*-driven CRISPR-Cas9 lentivirus-based approach. The inactivation of *Acly,* but not *Acss2*, ameliorated EAE (**Fig. 3a**, **Extended Data Fig. 5a)**, decreasing the expression of transcriptional modules linked to astrocyte epigenetic memory, inflammation, and NF-kB signaling (**Figs. 3b**-**d**, **Supp. Table 6**), but not affecting the number of CD4^+^ T cells in the CNS or the periphery **(Extended Data Figs. 5c-e)**. *Acly* inactivation in astrocytes also suppressed demyelination and decreased transcriptional programs linked to acetyl-CoA biosynthesis and H3K27 acetylation, compared to scramble control (**Fig. 3d**, **Extended Data Figs. 5a,b)**. Indeed, we detected decreased H3K27ac marks in *Il6* and *Ccl2* promoters following *Acly* inactivation (**Fig. 3e**). In addition, *Acly* inactivation in astrocytes down-regulated p300-driven NF-κB signaling during EAE (**Fig. 3f**, **Supp. Table 7)**. Finally, in support for a role of ACLY in p300 activation in astrocytes during CNS inflammation, we detected an increase in nuclear ACLY^+^p300^+^ astrocytes in spinal cord WM during EAE (**Fig. 3g**, **Extended Data Fig. 5f)**.

**Figure 3:**
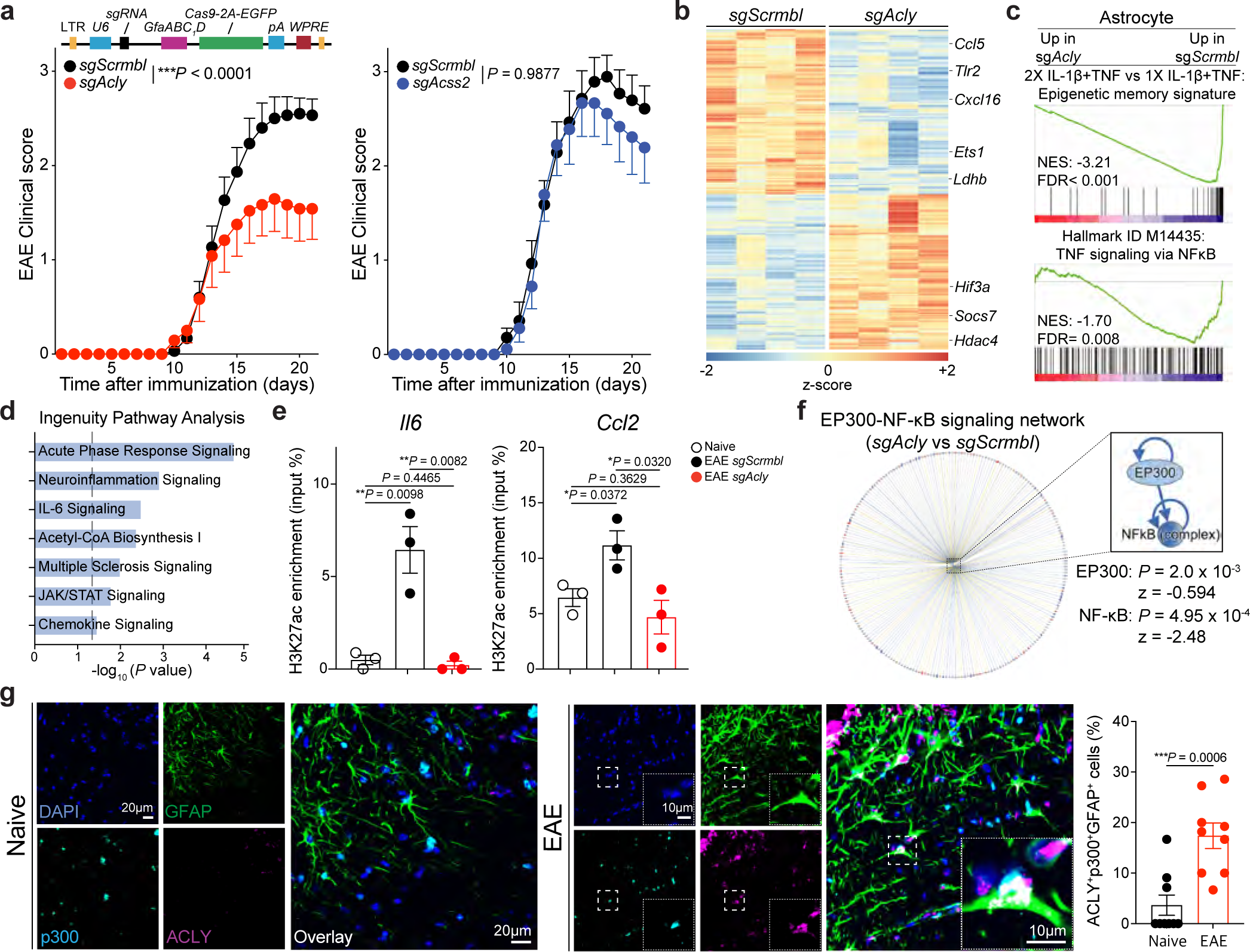
ACLY controls p300-derived astrocyte epigenetic memory in EAE. **(a)** EAE curves (sg*Scrmbl*; n=15; sg*Acly*; n=12; sg*Scrmbl*; n=14; sg*Acss2*; n=9). Representative data of at least two independent experiments. Two-way repeated measures of ANOVA. **(b)** Differential gene expression determined by RNA-seq in astrocytes from sg*Scrmbl*- and sg*Acly*-transduced mice 21 days after EAE induction (n=4 per group). **(c)** GSEA analysis comparing sg*Scrmbl*- and sg*Acly*-transduced astrocytes. **(d)** IPA pathway analysis down-regulated (blue) in comparing sg*Scrmbl*- and sg*Acly*-transduced astrocytes are shown. **(e)** Chip-qPCR analysis of the abundance of H3K27ac to pro-inflammatory gene promoters in comparing sg*Scrmbl*- and sg*Acly*-transduced astrocytes (n=3 per group). Unpaired two-sided *t*-test. **(f)** EP300-NF-κB signaling network comparing sg*Scrmbl*- and sg*Acly*-transduced astrocytes. **(g)** Immunostaining (left) and quantification (right) of ACLY^+^p300^+^ astrocytes in mice with/without EAE (n=9 spinal cord sections; n=3 mice per group). Unpaired two-sided *t*-test. Data shown as mean ± s.e.m.

To further validate these findings, we used the EAE model induced in NOD mice by MOG_35-55_ immunization, which recapitulates some aspects of secondary progressive MS^16,28^. First, we confirmed that IL-1β+TNF ICV administration in NOD EAE mice 124 days after disease induction (during the progressive disease phase) induces higher brain astrocyte *Il6*, *Ccl2*, and *Nos2* expression than in naïve mice **(Extended Data Figs. 6a-c)**. Moreover, we detected increased H3K27ac^+^p300^+^ and ACLY^+^p300^+^ astrocytes in spinal cord WM during NOD EAE **(Extended Data Figs. 6d,e)**. Finally, *Acly* or *Ep300* inactivation in CNS astrocytes using Gfap-driven CRISPR–Cas9 lentivirus suppressed NOD EAE progression, reduced demyelination, and decreased astrocyte transcriptional responses linked to H3K27 acetylation, epigenetic memory, NF-κB signaling, and inflammation compared to scramble control as determined by IHC and RNA-seq; no significant changes were detected in CD4^+^ T cells in the CNS or the periphery **(Extended Data Figs. 6f-i, 7a-e, Supp. Table 8)**. Taken together, these findings demonstrate that ACLY-p300-driven epigenetic memory in astrocytes promotes CNS pathology.

## FIND-seq analysis of Acly^+^Ep300^+^ astrocytes

Our studies identified a subset of ACLY/p300-controled astrocytes which promote EAE pathology. However, the intracellular localization of ACLY and p300 limits their use for the isolation and in-depth investigation of ACLY^+^p300^+^ astrocytes. To enable genome-wide transcriptional analyses of rare cells of interest isolated based on mRNA or DNA markers of interest, we recently described a novel method named focused interrogation of cells by nucleic acid detection and sequencing (FIND-seq)^29^. Thus, we used FIND-seq to analyze *Acly*^+^*Ep300*^+^ astrocytes.

First, we analyzed by FIND-seq *Acly*^-^*Ep300*^-^, *Acly*^+^*Ep300*^-^, *Acly*^-^*Ep300*^+^ and *Acly*^+^*Ep300*^+^ astrocytes isolated from the CNS of naïve and EAE TdTomato*^Gfap^* mice (**Figs. 4a,b**, **Extended Figs. 8a-f)**. In agreement with our previous IHC analyses, *Acly*^+^*Ep300*^+^ astrocytes were increased in EAE (**Fig. 4b**). Moreover, the transcriptional profile of *Acly*^+^*Ep300*^+^ astrocytes from EAE mice was closely correlated with the transcriptional signature of epigenetic memory defined based on the analysis of astrocytes following 2X IL-1β+TNF treatment *in vivo*; *Acly*^-^*Ep300*^+^, *Acly*^+^*Ep300*^-^ astrocyte subsets displayed intermediate profiles of this signature, suggesting that they represent transitional states towards the establishment of astrocyte epigenetic memory (**Fig. 4c**, **Extended Fig. 8g)**.

**Figure 4:**
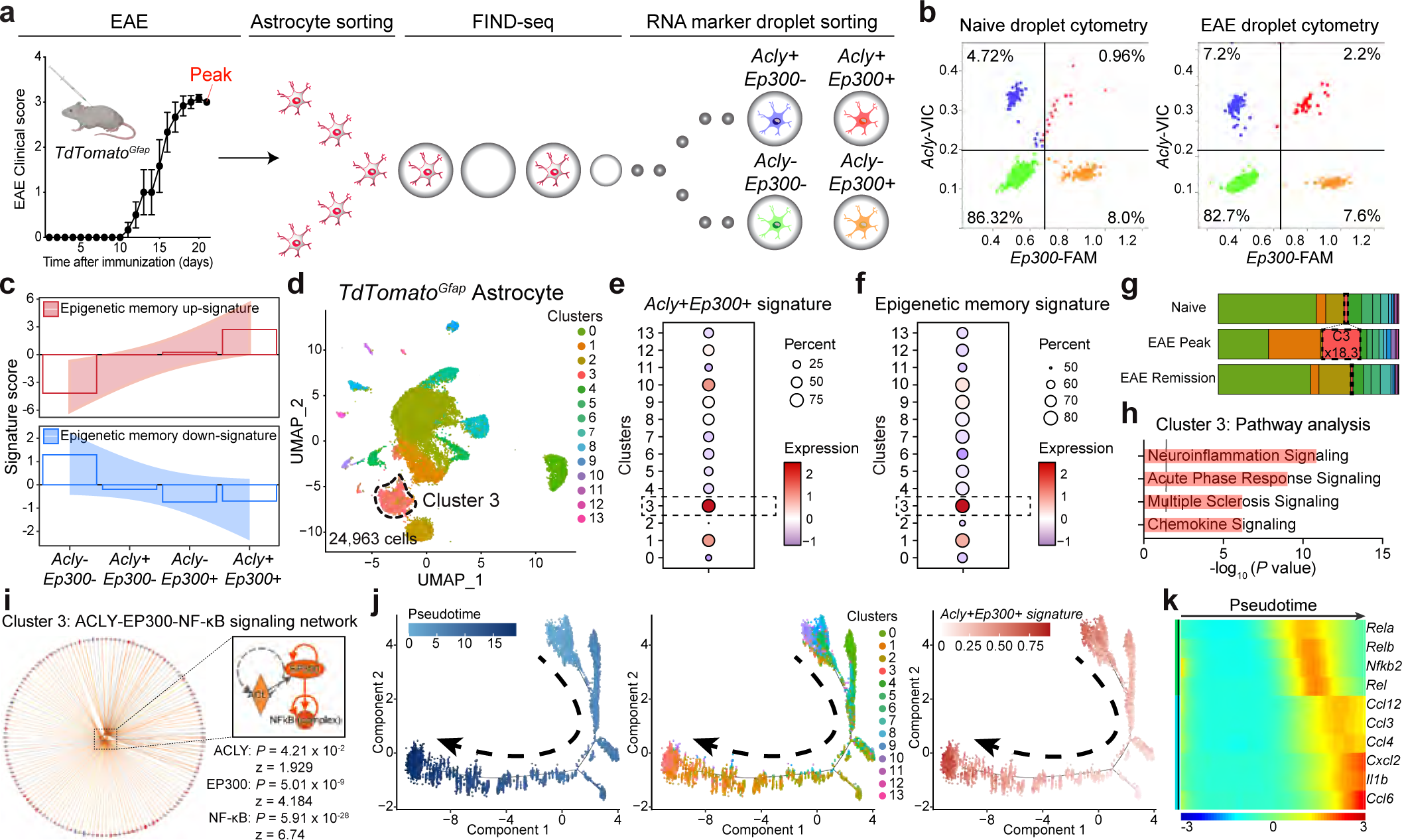
Analysis of *Acly^+^Ep300^+^*astrocytes with FIND-seq. **(a)** TdTomato*^Gfap^* astrocytes from naive and EAE mice were pooled (n=3 per group) and analyzed by FIND-seq using *Acly* and *Ep300* expression. EAE curve (EAE; n=3 per group). Two-way repeated measures of ANOVA. **(b)** Representative droplet cytometry plots. Numbers in each quadrant display the percentage of astrocytes in droplets. **(c)** Astrocyte epigenetic memory signature score in *Acly^-^Ep300^-^*, *Acly^+^Ep300^-^*, *Acly^-^Ep300^+^*, *Acly^+^Ep300^+^*EAE astrocytes. **(d)** UMAP plot of TdTomato*^Gfap^* astrocytes (n=24,963 cells)^10^. **(e)** *Acly^+^Ep300^+^*EAE astrocyte signature expression in EAE astrocyte clusters. **(f)** Astrocyte epigenetic memory signature score in EAE astrocyte clusters. **(g)** Fraction of cells per cluster overall. **(h)** IPA pathway analysis upregulated (red) in cluster 3 astrocytes. **(i)** ACLY-EP300-NF-κB signaling network of cluster 3 astrocytes. **(j)** Pseudotime trajectory in TdTomato*^Gfap^* astrocytes. Each cell is colored to indicate its pseudotime value **(k)** Pseudotime analysis of gene expression in TdTomato*^Gfap^* astrocytes.

Next, to interrogate the heterogeneity of memory astrocytes in the context of autoimmune CNS inflammation, we re-analyzed our astrocyte EAE scRNA-seq dataset generated in TdTomato*^Gfap^* mice^10^. Using the transcriptional signature of *Acly*^+^*Ep300*^+^ EAE astrocytes **(Supp. Table 9)**, we identified an astrocyte subset (cluster 3) that expanded 18.3-fold in EAE (**Figs. 4d-g**, **Extended Data Fig. 8h, Supp. Table 10)**. Indeed, the transcriptional signature of astrocytes in cluster 3 was the one most closely correlated with the transcriptional signature of astrocytes following 2X IL-1β+TNF treatment *in vivo* (**Fig. 4f**). Astrocytes in cluster 3 displayed the activation of pro-inflammatory pathways linked to astrocyte pathogenic activities and increased NF-κB signaling controlled by ACLY-p300-dependent epigenetic regulation (**Figs. 4h**,**i**, **Supp. Table 11)**. Moreover, we performed pseudotime analysis to interrogate the relationship across distinct reactive astrocyte subsets during EAE. Pseudotime analysis revealed a gradual transition toward cluster 3 astrocytes associated to epigenetic memory, which displayed a transcriptional profile enriched for genes linked to inflammation and NF-κB signaling (**Figs. 4j**,**k**). Interestingly, clusters 1 and 10 correspond to pathogenic astrocyte subsets expanded in EAE, which represent candidate intermediate states for the development of astrocyte pro-inflammatory memory controlled by p300-dependent NF-κB signaling (**Figs. 4e-g**, **Extended Data Figs. 8i-l)**. Thus, FIND-seq identified a disease-associated memory astrocyte subset controlled by ACLY- and p300-dependent epigenetic marks which promotes CNS inflammation.

## Astrocyte epigenetic memory in MS

Astrocyte subsets are thought to contribute to the pathology of MS^2,5^. Thus, we first investigated whether pro-inflammatory astrocyte epigenetic memory responses can be induced in human astrocytes. Human fetal astrocytes in culture displayed increased pro-inflammatory responses following 2X IL-1β+TNF stimulation, which were rescued by the treatment with the p300 inhibitor C646 treatment **(Extended Data Fig. 9a)**, resembling our findings in mouse astrocytes.

We next integrated two independent scRNA-seq datasets^30,31^, generating a dataset of 16,276 astrocytes from 17 MS patients and 12 controls. Unbiased clustering identified 6 astrocyte clusters in MS (**Fig. 5a**, **Extended Data Figs. 9b-e, Supp. Table 12,13)**. Consistent with our scRNA-seq studies in EAE (**Figs. 4d-i**), we identified an astrocyte subset (cluster 2) which displayed a transcriptional signature of similar to the one detected in mouse *Acly*^+^*Ep300*^+^ EAE astrocytes; this subset was expanded 24.5-fold in MS chronic lesions (20.4-fold in CA, 4.1-fold in CI) (**Figs. 5a-c**, **Extended Data Figs. 9f,g)**. This astrocyte subset displayed increased pro-inflammatory and NF-κB signaling (**Figs. 5d,e**). Indeed, IPA identified ACLY-p300 signaling as an upstream regulator of NF-κB signaling in cluster 2 astrocytes (**Figs. 5f**, **Supp. Table 14)**. Furthermore, pseudotime analysis revealed a transitional memory state towards cluster 2 astrocytes, which displayed a transcriptional profile enriched for genes related to histone acetylation, inflammation, and NF-κB signaling (**Figs. 5g,h**).

**Figure 5:**
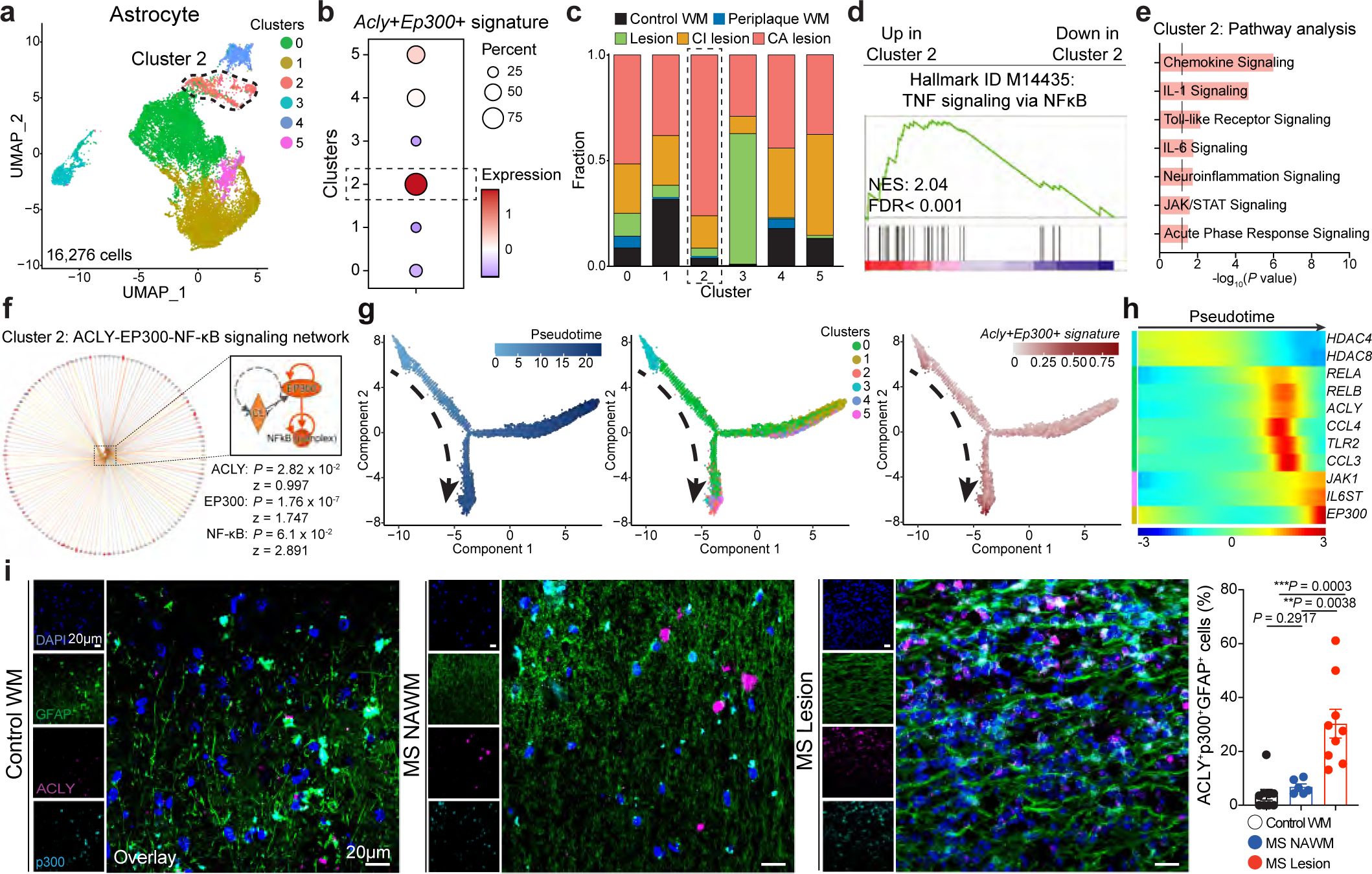
Identification of astrocyte epigenetic memory in MS. **(a)** UMAP plot of astrocytes from patients with MS and control individuals from Schirmer et al^31^. and Absinta et al^30^. (n=15,276 cells). **(b)** *Acly^+^Ep300^+^* EAE astrocyte signature expression in MS astrocyte clusters. **(c)** Cluster analysis of astrocytes based on percent composition in MS. WM, white matter; CI, chronic inactive; CA, chronic active. **(d)** GSEA analysis of cluster 2 astrocytes. **(e)** IPA pathway analysis up-regulated (red) in cluster 2 astrocytes. **(f)** ACLY-EP300-NF-κB signaling network of cluster 2 astrocytes. **(g)** Pseudotime trajectory in MS astrocytes. Each cell is colored to indicate its pseudotime value **(h)** Pseudotime analysis of gene expression in MS astrocytes. **(i)** Immunostaining and quantification of ACLY^+^, p300^+^, ACLY^+^p300^+^ astrocytes in tissue samples from patients with MS (n=9 sections (Lesion); n=6 sections (NAWM); n=3 per patient) and control individuals (n=3 sections; n=3 per patient). WM, white matter; NAWM, normally appearing white matter. Unpaired two-sided *t*-test. Data shown as mean ± s.e.m.

Finally, to validate these findings we analyzed CNS MS and control samples by IHC. In agreement with our scRNA-seq analyses, we detected an increase in nuclear ACLY^+^p300^+^ astrocytes in MS white matter lesions compared with control samples (**Fig. 5i**, **Extended Data Fig. 10a)**. Taken together, these data identify ACLY-p300 driven pro-inflammatory epigenetic memory astrocyte responses as potential contributors in CNS pathology during MS.

## Discussion

Astrocyte subsets defined by specific transcriptional activation states^6–8^ contribute to disease pathology in neurologic diseases^9–16^, but little is known about their stability. Here we describe ACLY^+^p300^+^ memory astrocytes which promote CNS pathology in EAE and, potentially, MS.

p300 and its paralog CBP regulate transcriptional programs via the acetylation of histones and other proteins^32^. The brain displays the highest p300/CBP HAT activity in the body^33^. Indeed, histone acetylation and other post-translational modifications, have been linked to long-term memory including recognition and contextual fear^33–36^. p300 also regulates cellular senescence and stress response, which promote systemic chronic inflammation^24,37–39^. However, little is known about the role of p300/CBP in CNS inflammation and neurodegeneration. Interestingly, a recent comparative study detected p300 upregulation in astrocytes in multiple pro-inflammatory contexts^40^, suggesting that p300-driven astrocyte epigenetic programs contribute to the pathogenesis of multiple neurologic conditions. In addition, p300 interacts with multiple partners including NF-κB^23^; a central regulator of pathogenic astrocyte responses^2,5^. Indeed, acetylation boosts the ability of NF-κB to drive pro-inflammatory transcriptional responses^41^. In this context, our findings suggest that p300-dependent histone acetylation promotes the activation of NF-kB-driven transcriptional modules, establishing an epigenetic memory in pro-inflammatory astrocytes. This increased activation of NF-kB-driven transcriptional modules may reflect both the increased accessibility of NF-kB to DNA responsive elements as well as the heightened recruitment of NF-kB via protein-protein interactions, boosting pathogenic activities in memory astrocytes.

Metabolic adaptations are integrated into epigenetic marks to control transcriptional responses^22,42^. Indeed, nuclear acetyl-CoA levels control HAT enzymatic activity^26^. Acetyl-CoA is generated by the metabolism of citrate or acetate catalyzed by ACLY or ACSS2^26^, respectively. For example, chromatin-recruited ACSS2 regulates CBP-mediated histone acetylation in hippocampal neurons during memory formation^36^. However, our studies suggest that the metabolic remodeling induced in astrocytes by proinflammatory cytokines^16^ is linked to ACLY-driven acetyl-CoA production, providing an example of how inflammation-induced metabolic adaptation in astrocytes can amplify astrocyte-driven pro-inflammatory responses. These findings resemble, previous observations on how inflammation-triggered changes in astrocyte sphingolipid metabolism boost NF-kB driven pro-inflammatory responses^16,28^.

ACLY is expressed by neurons and glial cells in the CNS^43^. ACLY catalyzes the synthesis of Acetyl-CoA, which is then used as an acetyl group donor by p300 during histone acetylation^27^. Interestingly, ACLY has been shown to participate in the control of T cell and macrophage responses^44–46^. These findings suggest that shared mechanisms operate in the regulation of the pro-inflammatory response of hematopoietic and non-hematopoietic cells such as astrocytes. Indeed, although we focused on astrocyte-intrinsic mechanisms of epigenetic memory, interactions with other cells in the CNS are also likely to modulate astrocyte epigenetic memory^2,5^. Microglia, for example, are important regulators of astrocyte responses^2,5^. Astrocytes also establish regulatory interactions with peripheral immune cells recruited to the CNS in homeostasis and pathology^10,11,14^. Finally, recent reports suggest that neurons modulate astrocyte responses via the regulation of histone serotonylation^47^. Hence, cell-cell interactions with CNS-resident and peripheral cells may integrate multiple stimuli to regulate astrocyte epigenetic memory.

Recent reports described months-to years-long epigenetic memory in innate immune cells and their progenitors^18,21,48,49^. Our findings suggest that astrocytes, non-hematopoietic CNS cells, also display epigenetic memory. Considering the long lifespan and low turnover rate of astrocytes^50,51^, these findings suggest that astrocyte subsets acquire epigenetic modifications which condition their long-term responses, with important implications for the pathology of chronic neurologic diseases. Hence, understanding the stability, heterogeneity and regulation of astrocyte epigenetic memory has the potential to provide new insights into the pathology of neurologic diseases and guide novel therapeutic approaches.

In summary, we identified a subset of ACLY^+^p300^+^ memory astrocytes controlled by epigenetic programs induced in the context of autoimmune CNS inflammation. These findings constitute an example of mechanisms of structural immunity^52^ driven by neuroimmune interactions that promote CNS pathology. Indeed, ACLY^+^p300^+^ memory astrocytes were most expanded in MS chronic active lesions, which have linked to MS progression^30^. In this context, recent developments in our molecular understanding of the mechanism of action of ACLY have guided the development of specific inhibitors, which are considered promising therapeutic agents for cancer and metabolic conditions^53–55^. Hence, ACLY inhibitors are candidate tools for the therapeutic modulation of ACLY^+^p300^+^ memory astrocytes in MS and other neurologic disorders.

## Figure Legends

**Extended Data Figure 1:**
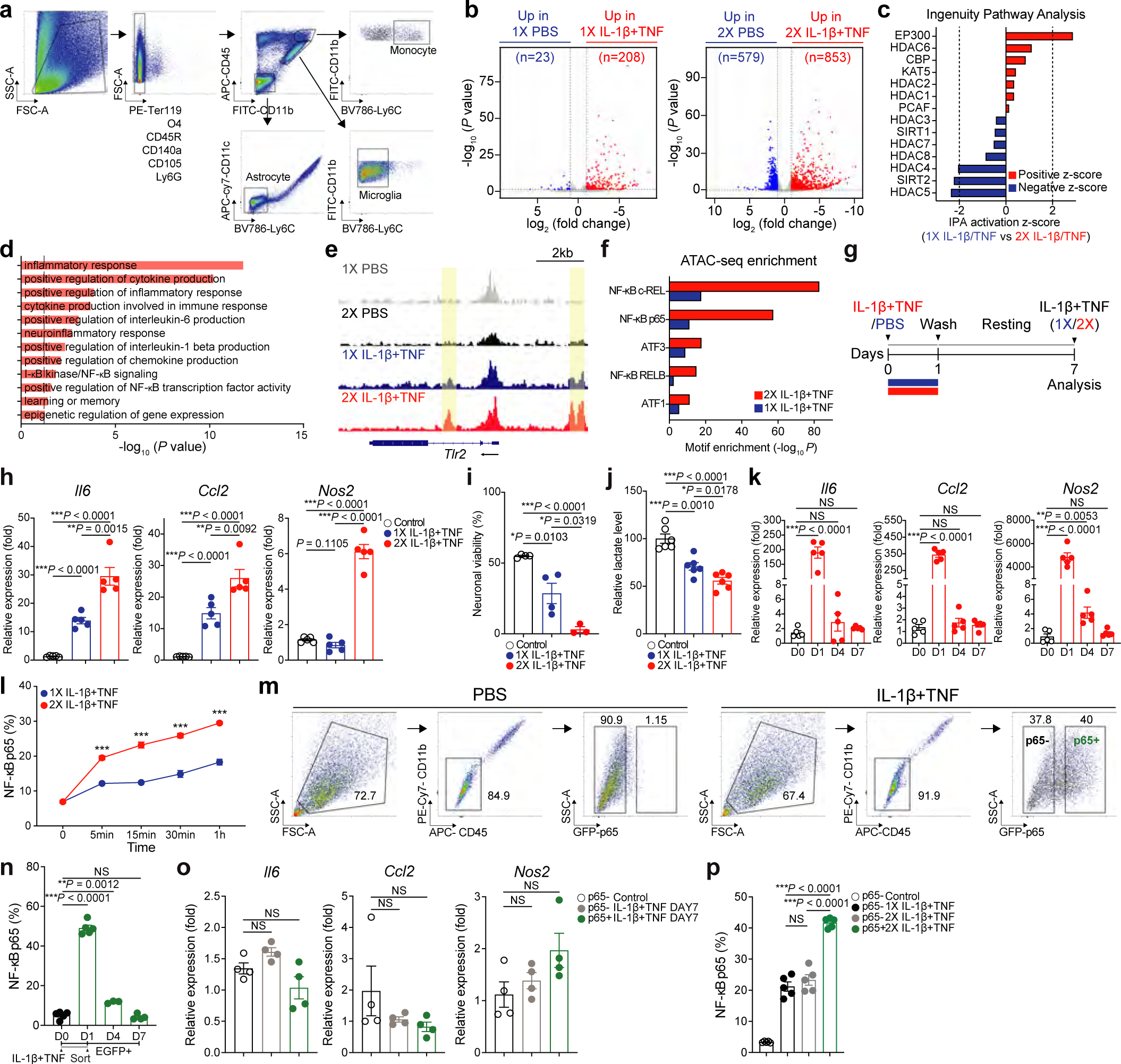
Analysis of astrocyte epigenetic memory *in vivo* and *in vitro*. **(a)** Fluorescence-activated cell sorting (FACS) sorting schematic for astrocytes, microglia, and monocytes. **(b)** Volcano plot of differential gene expression increased (red) or decreased (blue) analyzed by RNA-seq of sorted astrocytes. **(c)** IPA predicted upstream regulators in isolated astrocytes. Up-regulated (red) or down-regulated (blue) in astrocytes stimulated twice (2X IL-1β+TNF) versus once (1X IL-1β+TNF) are shown. **(d)** GO pathway analysis of ATAC-seq accessible peaks of isolated astrocytes comparing 2X IL-1β+TNF versus 1X IL-1β+TNF. **(e)** Genome browser snapshots showing the ATAC-seq sequencing tracks at the *Tlr2* locus. Only regions showing a significant increase (*p*-value < 0.05) in accessibility in astrocytes from mice stimulated twice (2X) versus once (1X) are highlighted by yellow boxes. **(f)** Homer DNA-motif enrichment analyses of differentially accessible peaks. **(g)** Experimental design for (h) to (j). Primary astrocytes received IL-1β/TNF stimulation once (1X) or twice (2X). **(h)** qPCR of astrocytes after 30min activation with IL-1β/TNF on day 7 (n=5 per group). Unpaired two-sided *t*-test. **(i)** Neuronal viability assay (n=4 control; n=4 1X; n=3 2X). Unpaired two-sided *t*-test. **(j)** Effect of IL-1β/TNF stimulation on lactate release (n=6 per group). Unpaired two-sided *t*-test. **(k)** qPCR analysis of astrocyte response after the first IL-1β/TNF stimulation (n=5 per group). Unpaired two-sided *t*-test. **(l)** FACS analysis of EGFP expression in IL-1β/TNF stimulated primary astrocytes isolated from *p65^EGFP^* reporter mice (0 min; n=5; Other time point; n=6 per group). \*\*\**P* < 0.0001. Unpaired two-sided *t*-test. **(m)** FACS sorting schematic of EGFP positive/negative astrocytes after PBS or IL-1β/TNF stimulation for 18-24h. **(n)** Primary astrocytes isolated from *p65^EGFP^* reporter mice received IL-1β/TNF stimulation. After 18-24h, EGFP positive astrocytes were sorted and analyzed by FACS of EGFP expression (n=3-5 per group). Unpaired two-sided *t*-test. **(o)** Primary astrocytes isolated from *p65^EGFP^* reporter mice received PBS or IL-1β/TNF stimulation. After 18-24h, EGFP positive/negative astrocytes were sorted and cultured for 6 days to perform qPCR (n=4 per group). Unpaired two-sided *t*-test. **(p)** p65 activation in EGFP positive/negative astrocytes 1h after IL-1β/TNF stimulation on day 7 (n=5 per group). Unpaired two-sided *t*-test. Data shown as mean ± s.e.m.

**Extended Data Figure 2:**
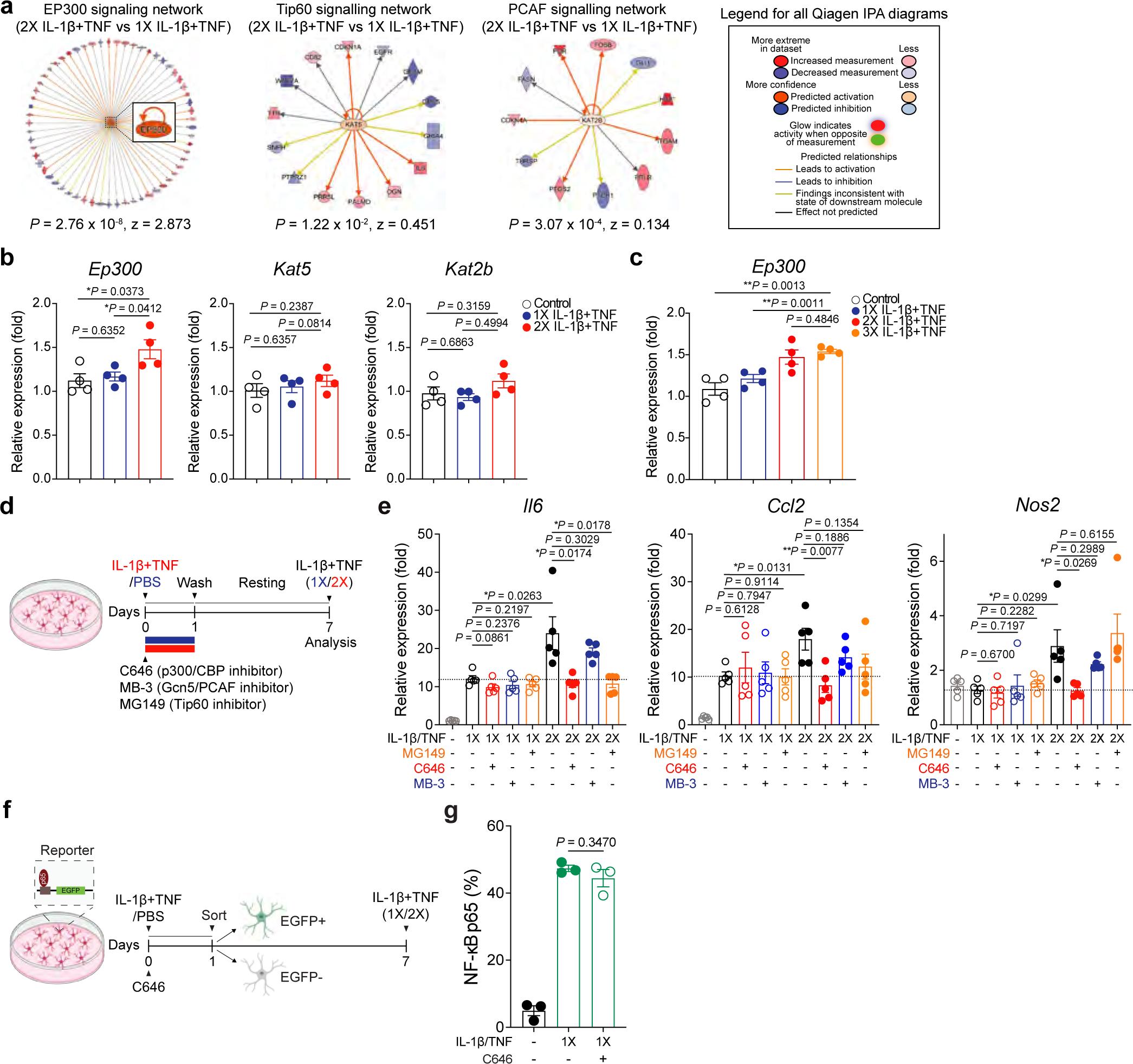
HAT enzyme expression and regulation of astrocyte epigenetic memory. **(a)** EP300, Tip60 (KAT5), and PCAF (KAT2B) signaling network in isolated brain astrocytes that received ICV administration of IL-1β/TNF twice (2X) compared to once (1X). **(b)** qPCR analysis of *Ep300*, *Kat5*, and *Kat2b* expression in primary astrocytes stimulated after 30min IL-1β/TNF stimulation on day 7 (n=4 per group). Unpaired two-sided *t*-test. **(c)** Primary astrocytes were stimulated with IL-1β/TNF once (1X), twice (2X), or three times (3X). qPCR analysis of astrocytes after 30min activation with IL-1β/TNF on day 14 (n=4 per group). Unpaired two-sided *t*-test. **(d)** Experimental design for (e). **(e)** qPCR of primary astrocytes in the presence with/without C646 (p300/CBP inhibitor), MB-3 (Gcn5/PCAF inhibitor), and MG149 (Tip60 inhibitor) after 30min stimulation with IL-1β/TNF on day 7 (n=5 per group). Unpaired two-sided *t*-test. **(f)** Experimental design for Fig 1j. **(g)** Gene expression 30min after IL-1β/TNF stimulation on day 7 of EGFP positive/negative astrocytes treated with C646 or vehicle (n=8 per group). Unpaired two-sided *t*-test. Data shown as mean ± s.e.m.

**Extended Data Figure 3:**
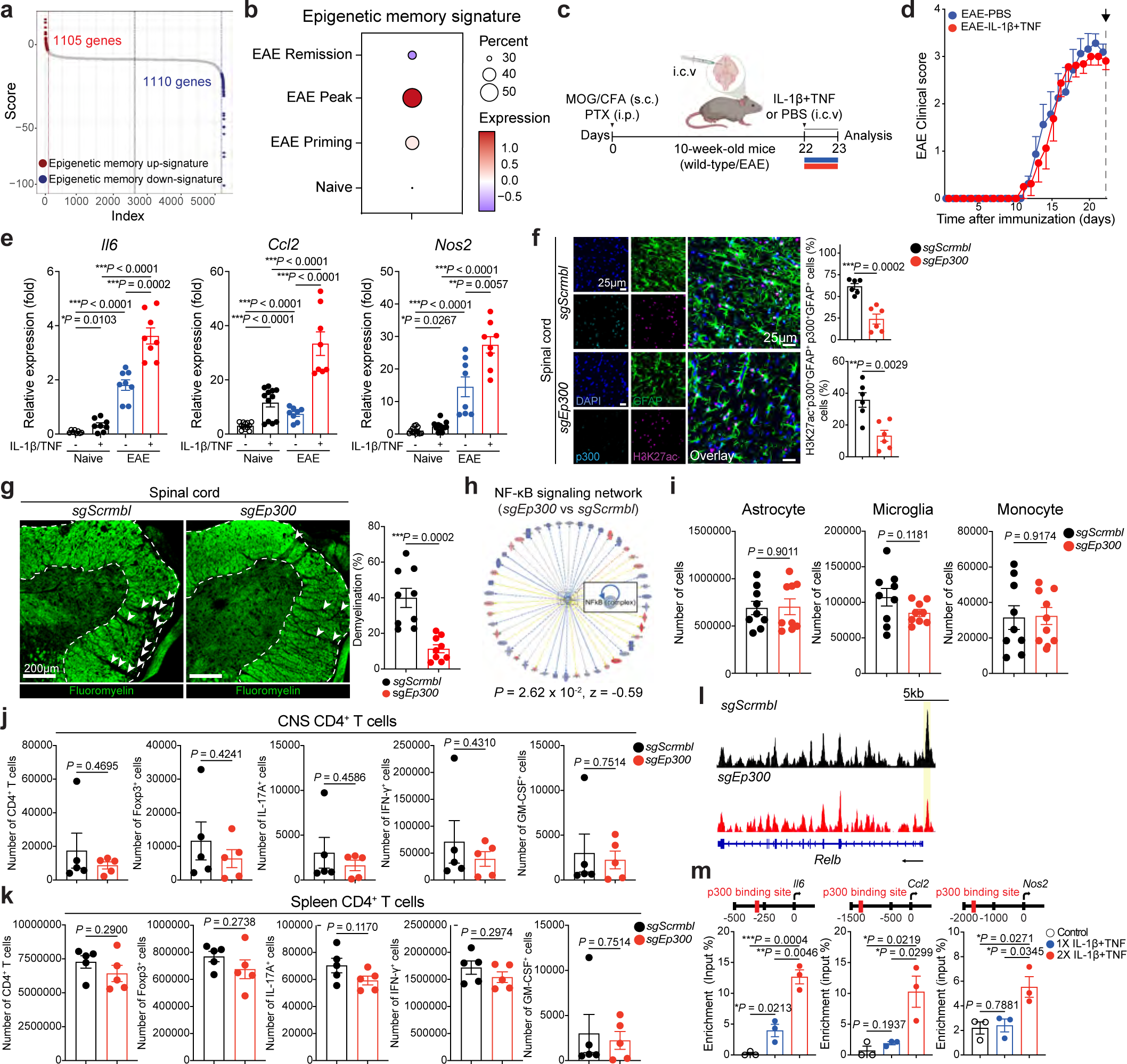
Astrocyte epigenetic memory and *Ep300* signaling in EAE. **(a)** Generation of astrocyte epigenetic memory signature filtered based on adjusted *p*-value and fold change. Up-signature (red) or down-signature (blue) in astrocytes stimulated twice (2X) than once (1X) are shown. **(b)** Astrocyte epigenetic memory signature score applied to naive and EAE scRNA-seq astrocyte dataset (Priming, Peak, and Remission)^10^. **(c)** Experimental design for (d,e). **(d)** EAE score for (e) (n=8 per group). Data shown as mean ± s.e.m. Naïve and EAE induced C57BL/6 mice received ICV administration of IL-1β/TNF (EAE peak, Day 22), and 18-24h later sorted brain astrocytes were analyzed. **(e)** qPCR of IL-1β/TNF response of astrocytes (Naïve; n=12; EAE; n=8 per group). Representative data of two independent experiments. Unpaired two-sided *t*-test. **(f)** Immunostaining (left) and quantification (right) of H3K27ac^+^ and p300^+^ astrocytes from sg*Scrmbl*- and sg*Ep300*-transduced mice at 23 days after EAE induction (n=6 spinal cord sections; n=3 mice per group). Unpaired two-sided *t*-test. **(g)** FluoroMyelin dye staining and percentage of myelin loss in spinal cord from sg*Scrmbl*- and sg*Ep300*-treated mice (n=9 spinal cord sections; n=3 mice per group). Lesions indicated by arrowheads. Unpaired two-sided *t*-test. **(h)** NF-kB signaling network comparing sg*Ep300*-transduced versus sg*Scrmbl-*transduced astrocytes. **(i)** Quantification of CNS-resident cells from sg*Scrmbl*- and sg*Ep300*-transduced mice (n=9 per group). Unpaired two-sided *t*-test. **(j,k)** Analysis of CNS T cells (up) and splenic T cells (bottom) from sg*Scrmbl*- or sg*Ep300*-transduced mice (n=5 per group). Unpaired two-sided *t*-test. **(l)** Genome browser snapshots showing the *Relb* locus. Only regions showing a significant decrease (*p*-value < 0.05) in accessibility in sg*Scrmbl*-transduced versus sg*Ep300*-transduced astrocytes are highlighted by yellow boxes. **(m)** Chip-qPCR analysis of p300 recruitment to promoters in primary astrocytes 30min after IL-1β/TNF stimulation on day 7 (n=3 per group). Unpaired two-sided *t*-test. Data shown as mean ± s.e.m.

**Extended Data Figure 4:**
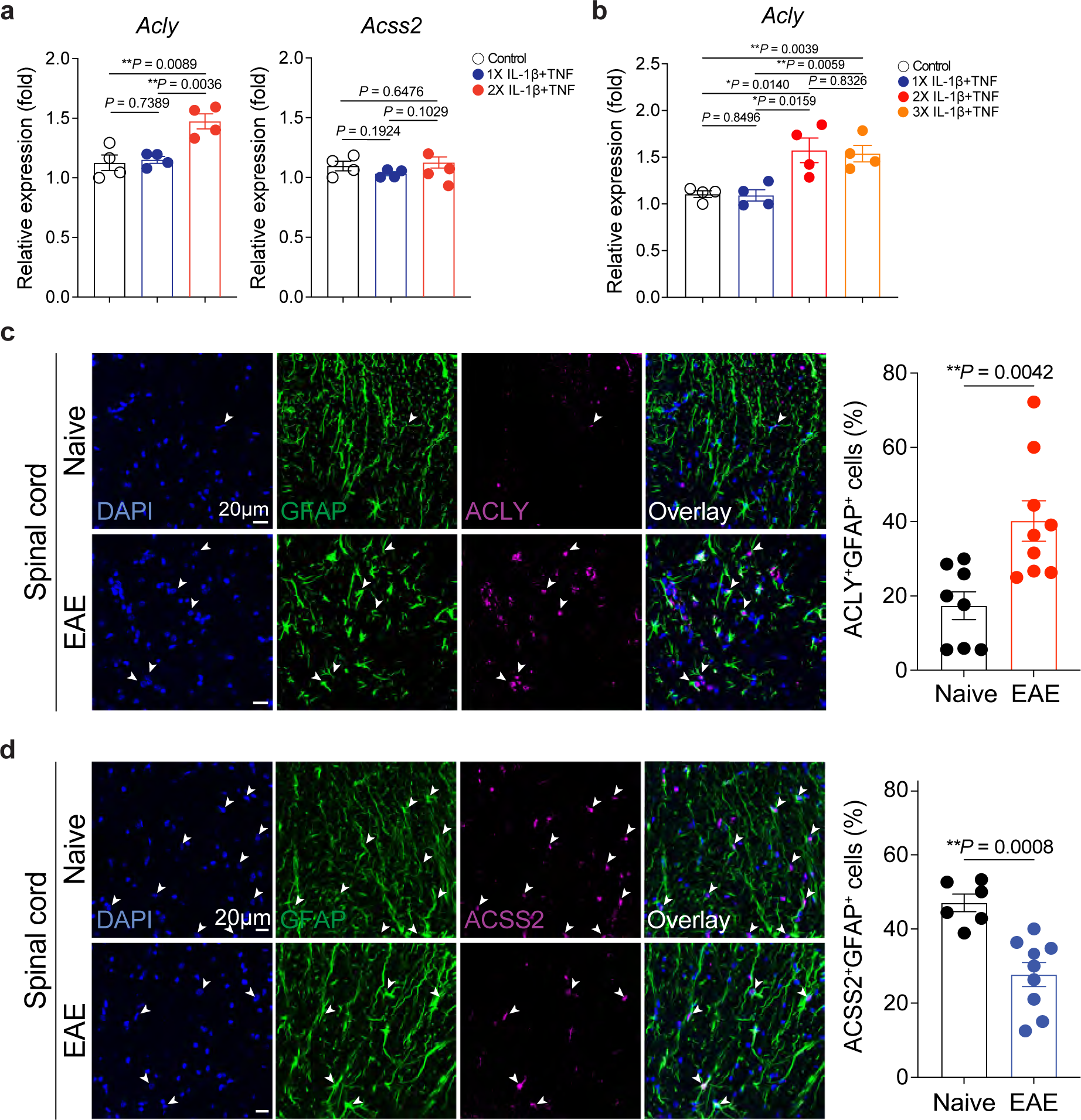
ACLY and ACSS2 signaling in EAE astrocytes. **(a)** qPCR analysis of *Acly* and *Acss2* expression in primary astrocytes stimulated for 30min with IL-1β/TNF on day 7 (n=4 per group). Unpaired two-sided *t*-test. **(b)** Primary astrocytes received IL-1β/TNF stimulation once (1X), twice (2X), or three times (3X). qPCR analysis of astrocytes after 30min activation with IL-1β/TNF on day 14 (n=4 per group). Unpaired two-sided *t*-test. **(c)** Immunostaining (left) and quantification (right) of ACLY^+^ astrocytes in mice with/without EAE (n=8 spinal cord sections (naïve); n=9 spinal cord sections (EAE); n=3 mice per group). Unpaired two-sided *t*-test. **(d)** Immunostaining (left) and quantification (right) of ACSS2^+^ astrocytes in mice with/without EAE (n=6 spinal cord sections (naïve); n=9 spinal cord sections (EAE); n=3 mice per group). Unpaired two-sided *t*-test. Data shown as mean ± s.e.m.

**Extended Data Figure 5:**
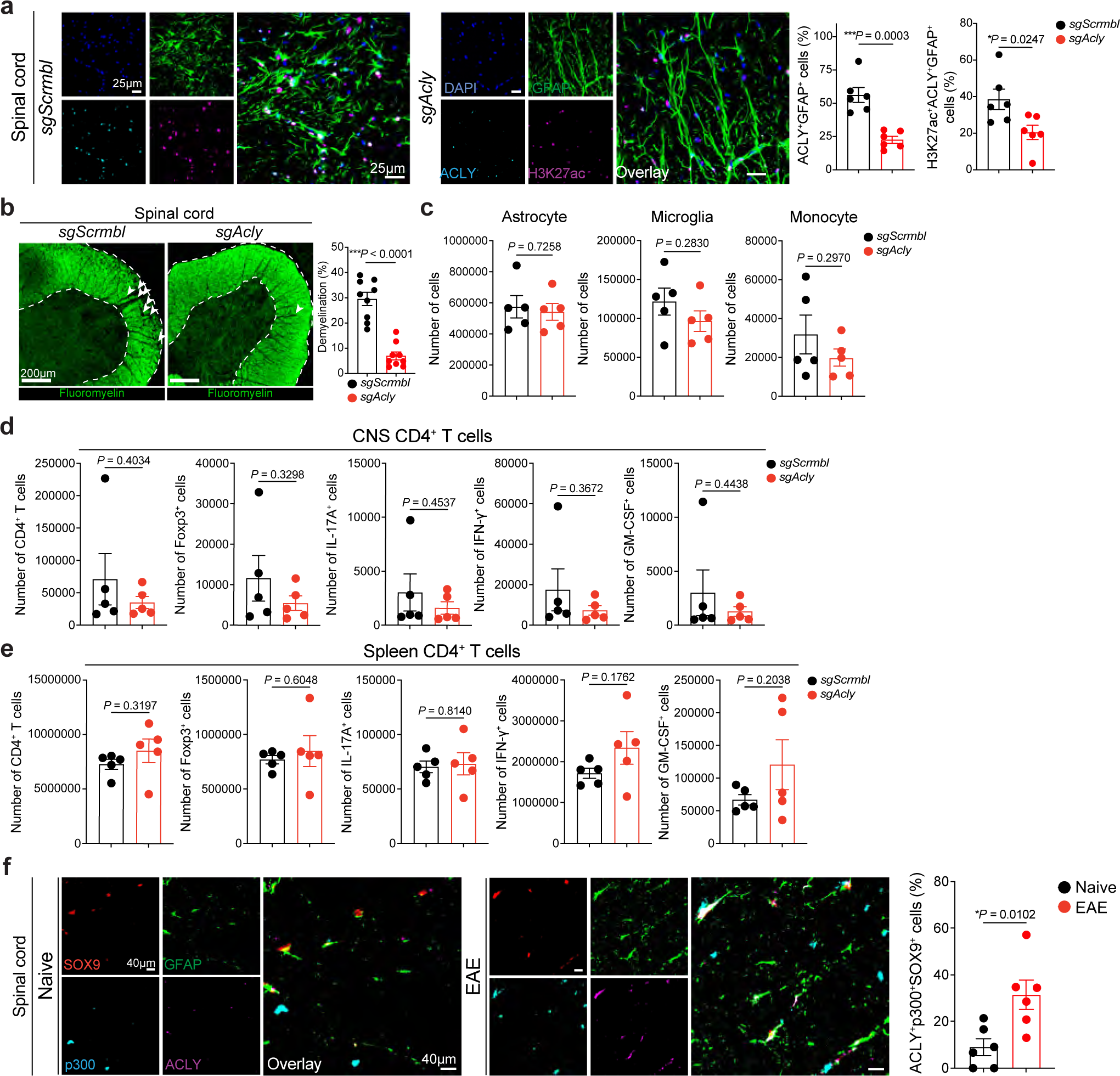
Astrocyte *Acly* signaling and ACLY+p300+ astrocytes in EAE. **(a)** Immunostaining (left) and quantification (right) of H3K27ac^+^ and ACLY^+^ astrocytes from sg*Scrmbl*- and sg*Acly*-transduced mice at 21days after EAE induction (n=6 spinal cord sections; n=3 mice per group). Unpaired two-sided *t*-test. **(b)** Staining with FluoroMyelin dye and percentage of myelin loss in spinal cord of sg*Scrmbl*- and sg*Acly*-treated mice (n=9 spinal cord sections; n=3 mice per group). Lesions indicated by arrowheads. Unpaired two-sided *t*-test. **(c)** Quantification of CNS-resident cells from sg*Scrmbl*- and sg*Acly*-transduced mice (n=5 per group). Unpaired two-sided *t*-test. **(d,e)** Analysis of CNS T cells (up) and splenic T cells (bottom) from sg*Scrmbl*- or sg*Acly*-transduced mice (n=5 per group). Unpaired two-sided *t*-test. Data shown as mean ± s.e.m. **(f)** Immunostaining (left) and quantification (right) of ACLY^+^SOX9^+^, p300^+^SOX9^+^, and ACLY^+^p300^+^SOX9^+^ astrocytes in EAE and control mice (n=6 spinal cord sections; n=3 mice per group). Unpaired two-sided *t*-test. Data shown as mean ± s.e.m.

**Extended Data Figure 6:**
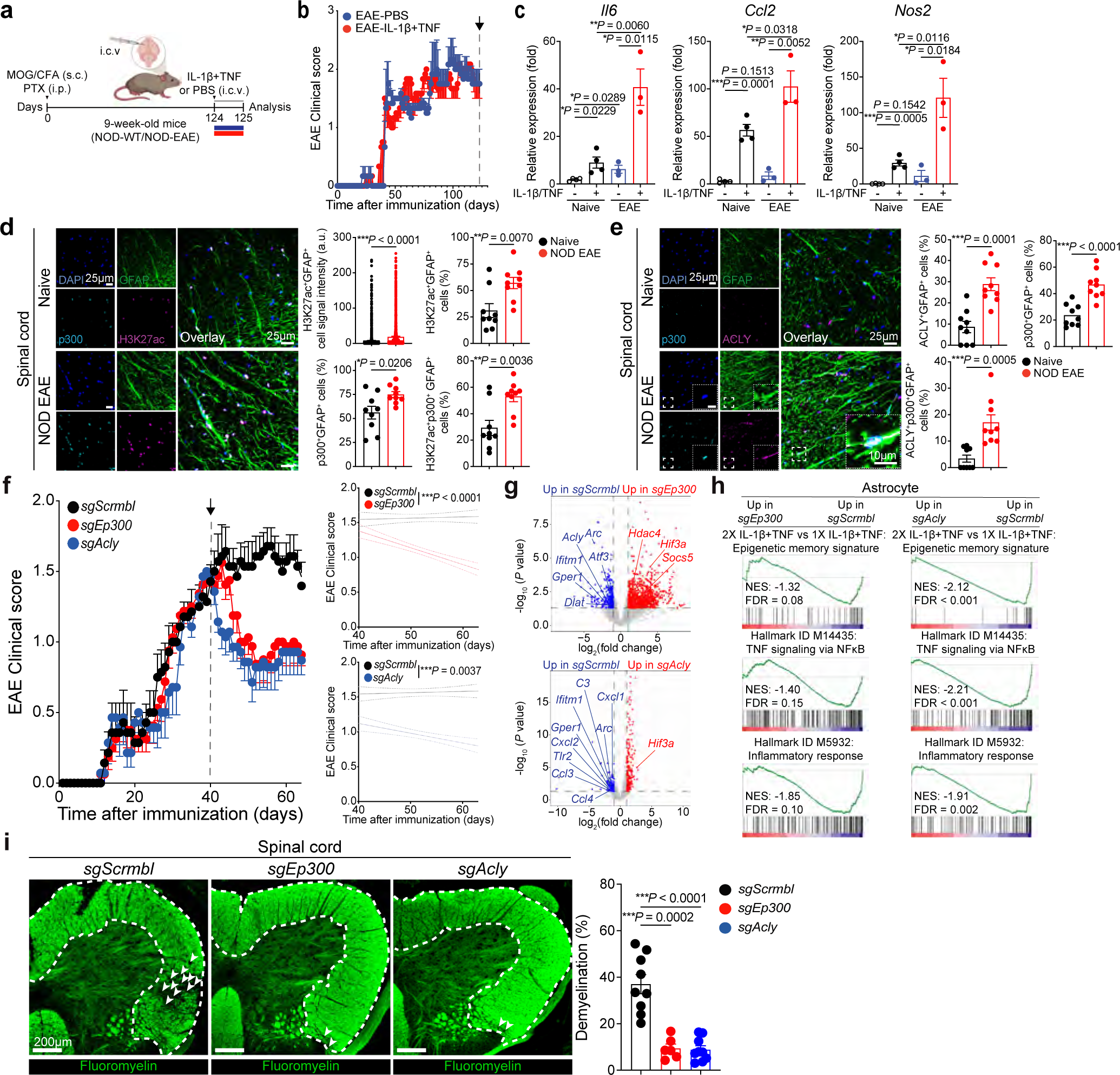
Astrocyte epigenetic memory in NOD EAE. **(a)** Experimental design for (c). **(b)** NOD EAE score for (c) (n=3 per group). **(c)** Naïve and EAE induced NOD mice received ICV administration of IL-1β/TNF (EAE progressive, Day 124). After 18-24h, sorted brain astrocytes were analyzed by qPCR (n=3 per group). Unpaired two-sided *t*-test. **(d)** Immunostaining (left) and quantification (right) of H3K27ac^+^ and p300^+^ astrocytes in mice with/without NOD EAE (n=9 spinal cord sections; n=3 mice per group). Astrocyte H3K27ac levels were calculated as the mean signal intensity (arbitrary units) per GFAP+ cells using automated unbiased quantification. Unpaired two-sided *t*-test. **(e)** Immunostaining (left) and quantification (right) of ACLY^+^, p300^+^, and ACLY^+^p300^+^ astrocytes in mice with/without NOD EAE (n=9 spinal cord sections; n=3 mice per group). Unpaired two-sided *t*-test. **(f)** NOD EAE curves (sg*Scrmbl*; n=7; sg*Ep300*; n=8; sg*Acly*; n=7). Lentivirus were injected at day 40. Representative data of two independent experiments. Regression slope two-sided *t*-test compared with sg*Scrmbl*. **(g)** Volcano plot of differential gene expression determined by RNA-seq in astrocytes isolated from sg*Scrmbl*-, sg*Ep300*-, and sg*Acly*-transduced mice 64 days after NOD EAE induction (n=3 sg*Scrmbl*, n=3 sg*Ep300*, n=2 sg*Acly*). **(h)** GSEA analysis comparing sg*Scrmbl*-, sg*Ep300*-, and sg*Acly*-transduced astrocytes. **(i)** Staining with FluoroMyelin dye and percentage of myelin loss from sg*Scrmbl*-, sg*Ep300*-, and sg*Acly*-transduced mice spinal cord (n=6 spinal cord sections (*sgEp300*); n=9 spinal cord sections (*sgScrmbl, sgAcly*); n=3 mice per group. Lesions indicated by arrowheads. Unpaired two-sided *t*-test. Data shown as mean ± s.e.m.

**Extended Data Figure 7:**
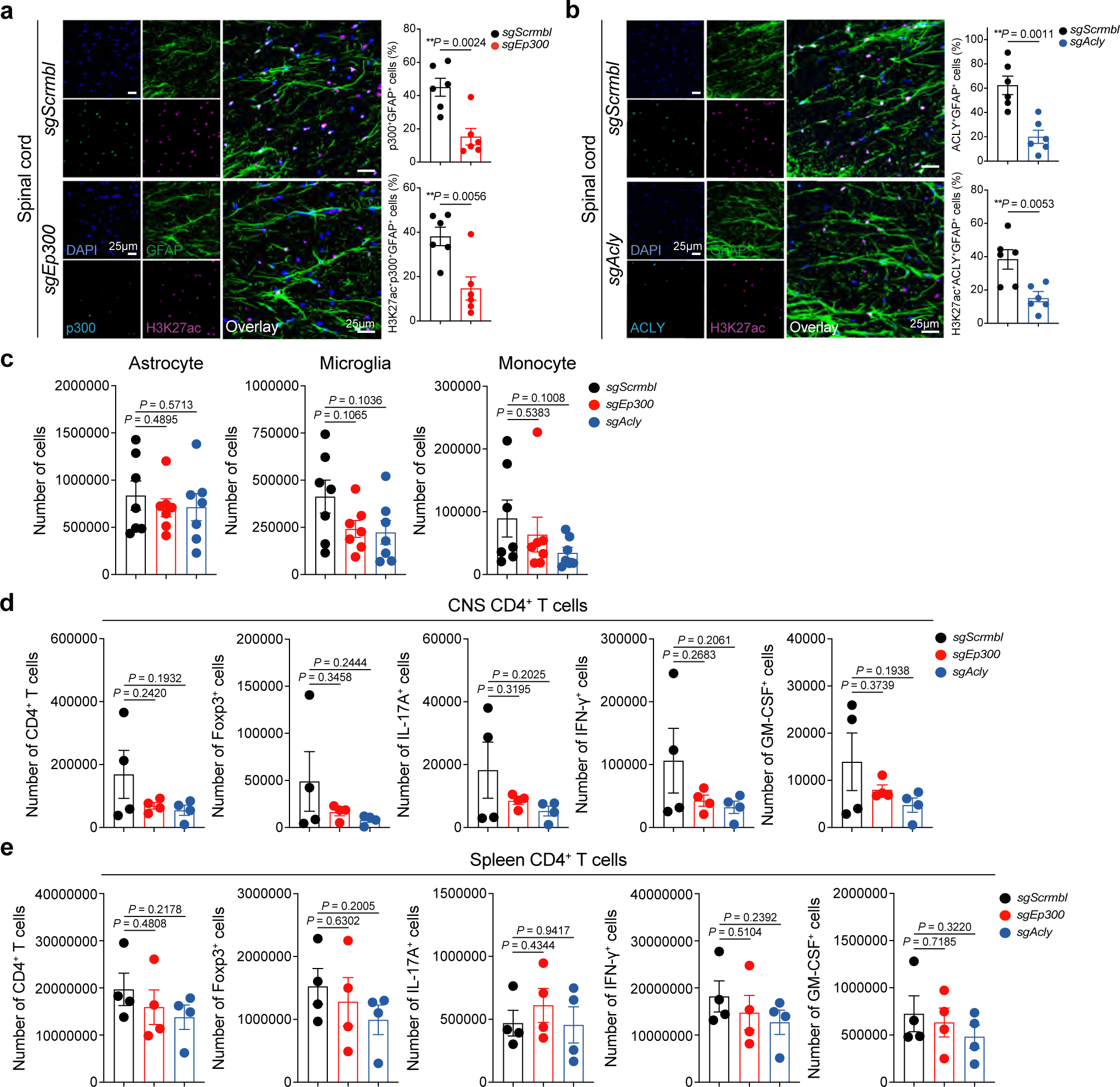
ACLY and p300 signaling in NOD EAE astrocytes. **(a)** Immunostaining (left) and quantification (right) of H3K27ac^+^ and p300^+^ astrocytes from sg*Scrmbl*- and sg*Ep300*-transduced NOD mice 64 days after EAE induction (n=6 spinal cord sections; n=3 mice per group). Unpaired two-sided *t*-test. **(b)** Immunostaining (left) and quantification (right) of H3K27ac^+^ and ACLY^+^ astrocytes from sg*Scrmbl*- and sg*Acly*-transduced NOD mice 64 days after EAE induction (n=6 spinal cord sections; n=3 mice per group). Unpaired two-sided *t*-test. **(c)** Quantification of CNS-resident cells from sg*Scrmbl*-, sg*Ep300*, and sg*Acly*-transduced mice (n=7 per group). Unpaired two-sided *t*-test. **(d,e)** Analysis of CNS T cells (up) and splenic T cells (bottom) from sg*Scrmbl*-, sg*Ep300*, and sg*Acly*-transduced mice (n=4 per group). Unpaired two-sided *t*-test. Data shown as mean ± s.e.m.

**Extended Data Figure 8:**
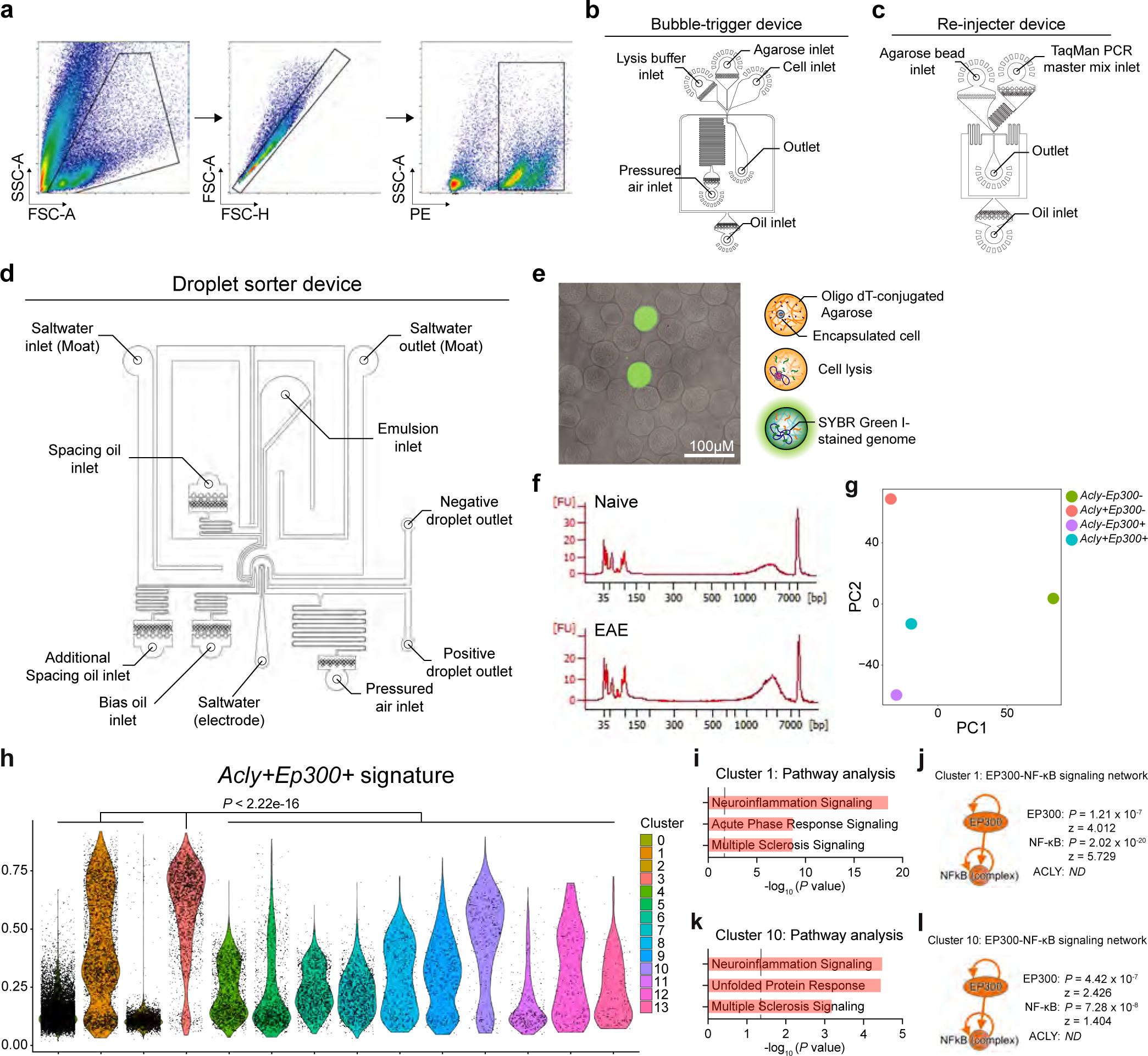
Analysis of *Acly^+^Ep300^+^* astrocytes by FIND-seq. **(a)** FACS sorting schematic for tdTomato*^Gfap^* astrocytes. **(b-d)** Schematic illustration of microfluidic devices utilized in FIND-seq. **(b)** The bubble-triggered device has with five inlets: i) cell inlet, ii) Oligo dT primer-conjugated agarose inlet, iii) lysis buffer inlet, iv) oil inlet, and v) pressurized air inlet. **(c)** The re-injector device has three inlets: i) agarose bead inlet, ii) TaqMan PCR master mix inlet, and iii) oil inlet. **(d)** The droplet sorter device has seven inlets: i) emulsion inlet, ii) spacing oil inlet, iii) additional spacing oil inlet, iv) bias oil inlet, v) saltwater inlet (for the electrode), vi) saltwater inlet (for the moat), and vii) pressurized air inlet. **(e)** Astrocytes are encapsulated in an agarose bead along with the lysis buffer. The genome entrapped in the agarose bead is stained with SYBR Green I and visualized using a fluorescence microscope. **(f)** cDNA, produced on the agarose bead, is amplified via WTA and validated using a Bioanalyzer. **(g)** Principal component analysis (PCA) plot of *Acly^-^ Ep300^-^*, *Acly^+^Ep300^-^*, *Acly^-^Ep300^+^*, *Acly^+^Ep300^+^* EAE astrocytes. **(h)** Violin plot depicting *Acly^+^Ep300^+^* EAE astrocyte signature expression in EAE astrocytes. **(i)** IPA pathway analysis up-regulated (red) in cluster 1 astrocytes. **(j)** EP300-NF-κB signaling network of cluster 1 astrocytes. **(k)** IPA pathway analysis up-regulated (red) in cluster 10 astrocytes. **(l)** EP300-NF-κB signaling network of cluster 10 astrocytes.

**Extended Data Figure 9:**
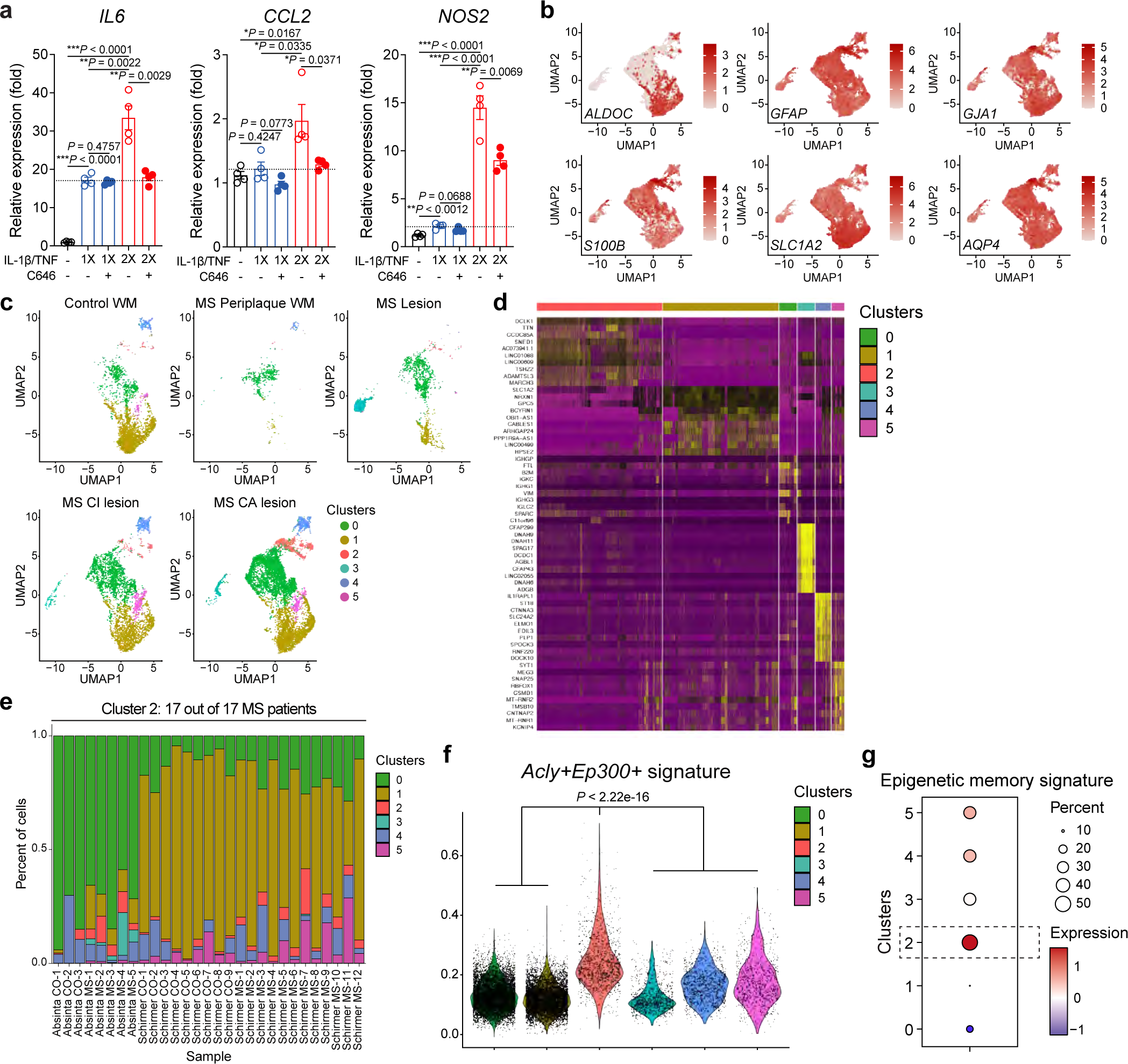
Analysis of human astrocyte epigenetic memory and MS astrocyte scRNA-seq. **(a)** Primary human fetal astrocytes received IL-1β/TNF stimulation once (1X) or twice (2X). qPCR of astrocytes in the presence with/without C646 (p300/CBP inhibitor) after 2h stimulation with IL-1β/TNF on day 7 (n=4 per group). Unpaired two-sided *t*-test. **(b)** Gene scatterplots of astrocyte markers. **(c)** Unsupervised clustering UMAP plot of astrocytes from patients with MS and control individuals from Schirmer et al^31^. and Absinta et al^30^. (n=16,276 cells). WM, white matter; CI, chronic inactive; CA, chronic active. **(d)** Significantly enriched genes by astrocyte cell type cluster. **(e)** Cluster distribution of CNS cells. **(f)** Violin plot depicting *Acly^+^Ep300^+^*EAE astrocyte signature expression in MS astrocytes. **(g)** Astrocyte epigenetic memory signature score in astrocyte clusters of control and MS patients.

**Extended Data Figure 10:**
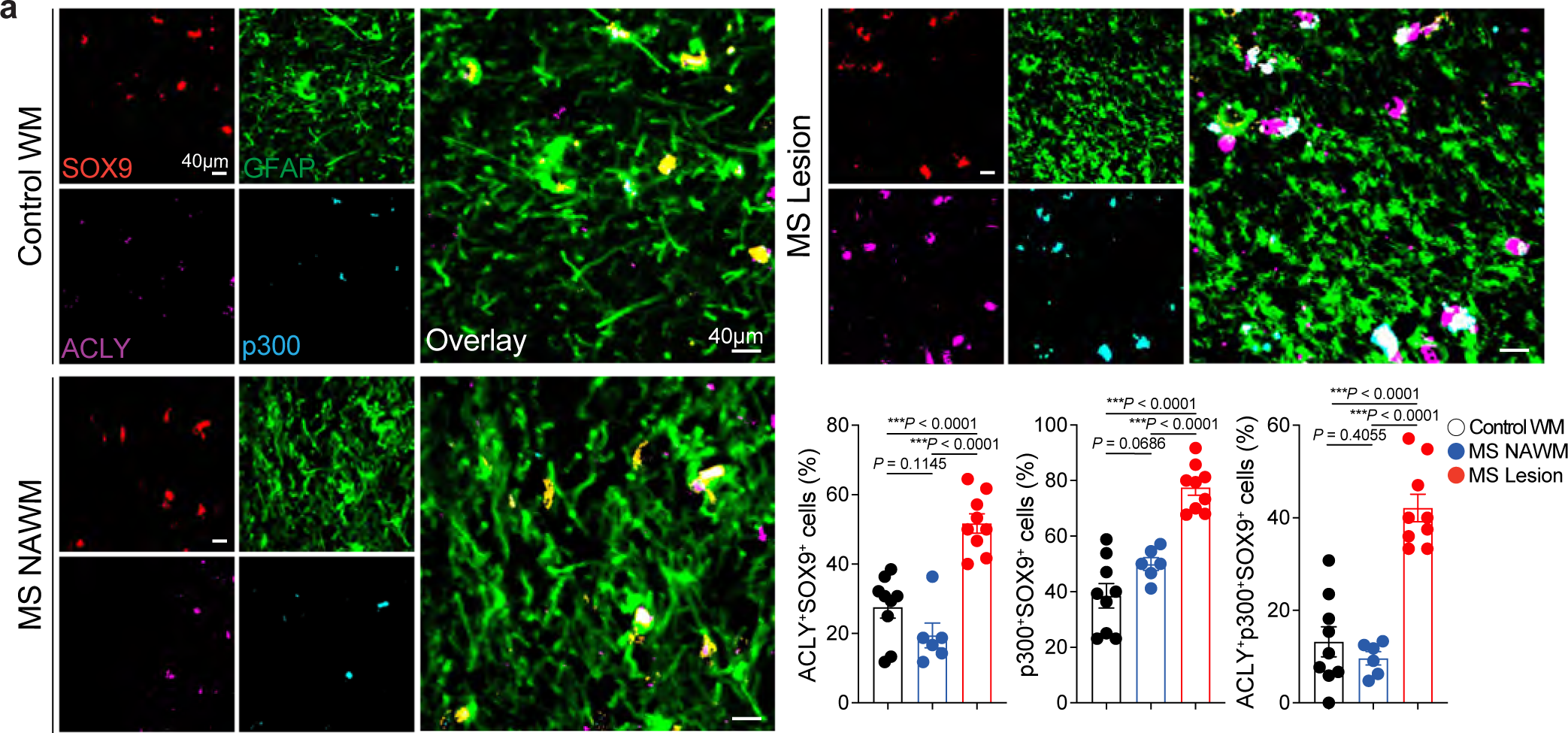
ACLY+p300+ astrocytes in MS patients. **(a)** Immunostaining and quantification of ACLY^+^SOX9^+^, p300^+^SOX9^+^, ACLY^+^p300^+^SOX9^+^ astrocytes in CNS samples from MS patients with MS (n=9 sections (Lesion); n=6 sections (NAWM); n=3 per patient) and controls (n=3 sections; n=3 per patient). WM, white matter; NAWM, normally appearing white matter. Unpaired two-sided *t*-test. Data shown as mean ± s.e.m.

## Methods

### Mice

Adult male and female mice, and postnatal pups were used on a C57BL/6J background (#000664, The Jackson Laboratory); NOD/ShiLtJ (#001976, The Jackson Laboratory) mice were obtained from the Jackson Laboratory. B6.Cg-Tg(Gfap-cre)73.12Mvs/J mice^56^ (The Jackson Laboratory, #012886) mice were crossed with B6.Cg-Gt(ROSA)26Sortm9(CAG-tdTomato)Hze/J mice^57^ (The Jackson Laboratory, #007909) to generate *Gfap*(*Cre/+*)*;TdTomato*(*f/+*) (*TdTomato^Gfap^*) mice. Postnatal pups were used from the NF-kB reporter strain FVB.Cg-Tg(HIV-EGFP,luc)8Tsb/J^58^ (The Jackson Laboratory, #027529) at P0-P3. Mice were kept in a pathogen-free facility at the Hale Building for Transformative Medicine at Brigham and Women’s Hospital in accordance with the IACUC guidelines. 9–12-week-old mice were used for stereotactic injection and EAE induction. Pups were sacrificed between P0-P3 for harvesting and culturing astrocytes. All procedures were reviewed and approved under the IACUC guidelines at Brigham and Women’s Hospital.

### In vivo cytokine stimulation

C57BL/6J mice at age 10-12 weeks were anesthetized using 1-3% isoflurane mixed with oxygen. Heads were shaved and cleaned using 70% ethanol and Betadine (Thermo Fisher, #19-027132) followed by a medial incision of the skin to expose the skull. Mice were injected with 100 ng/mL IL-1β (R&D Systems, #401-ML-005, 100 µg/mL stock in PBS) and 50 ng/mL TNF (R&D Systems, #410-MT-010, 100 µg/mL stock in PBS) diluted in PBS or PBS. At day 7, mice were re-injected (2X) or injected (1X) with 100 ng/mL IL-1β and 50 ng/mL TNF or PBS for 18-24 hours and astrocytes were sorted for further analysis. For EAE experiments, mice were injected with 100 ng/mL IL-1β and 50 ng/mL TNF or PBS for 18-24 hours and astrocytes were sorted for further analysis. The lateral ventricles were targeted bilaterally using the coordinates: +/-1.0 (lateral), -0.44 (posterior), -2.2 (ventral) relative to Bregma. Mice were injected by two 5µL injections using a 25 µL Hamilton syringe (Sigma-Aldrich, #20787) on a stereotaxic alignment system (Kopf, #1900) and the incision was sutured.

### Primary astrocyte cultures from neonatal mice

Procedures were performed as described previously^59^. Brains of mice aged P0-P3 were dissected into PBS on ice. Cortices were discarded and the brain parenchyma were pooled, centrifuged at 500g for 10 minutes at 4°C and resuspended in 0.25% Trypsin-EDTA (Thermo Fisher Scientific, #25200-072) at 37°C for 10 minutes. Trypsin was neutralized by adding DMEM (Thermo Fisher Scientific, # 11965118) supplemented with 10% FBS (Thermo Fisher Scientific, #10438026) and 1% penicillin/streptomycin (Thermo Fisher Scientific, #15140148), and cells were passed through a 70µm cell strainer. Cells were centrifuged at 500g for 10 minutes at 4°C, resuspended in DMEM with 10% FBS/1% penicillin/streptomycin and cultured in T-75 flasks (Falcon, #353136) at 37°C in a humidified incubator with 5% CO2, for 7-10 days until confluency was reached. Astrocytes were shaken for 30 minutes at 180 rpm, the supernatant was collected for microglia and the media was changed, then astrocytes were shaken for at least 2 hours at 220 rpm and the supernatant was aspirated and the media was changed again. Medium was replaced every 2–3 days. Primary astrocytes were treated with 100 ng/mL IL-1β (R&D Systems, #401-ML-005, 100 µg/mL stock in PBS) and 50 ng/mL TNF (R&D Systems, #410-MT-010, 100 µg/mL stock in PBS) diluted in DMEM (Thermo Fisher Scientific, # 11965118) that was supplemented with 10% FBS (Life Technologies, #10438026) and 1% penicillin/streptomycin (Life Technologies, #15140122) or PBS. The medium was replaced and washed twice after 18-24 hours, then astrocytes were rested for 6 days. Medium was replaced every 2–3 days. At day 7, astrocytes were re-treated (2X) or treated (1X) with 100 ng/mL IL-1β and 50 ng/mL TNF or PBS for 30min. Astrocytes were stimulated and collected on day 14 for 3X treatment. For p65 sorting experiments, EGFP^pos^ and EGFP^neg^ astrocytes were FACS sorted after 18-24 hours of IL-1β and TNF treatment and re-seeded and rested for 6 days. Astrocytes were subsequently gated on: CD11b^neg^CD45^neg^EGFP^pos/neg^. Medium was replaced every 2–3 days. At day 7, astrocytes were re-treated (2X) or treated (1X) with 100 ng/mL IL-1β and 50 ng/mL TNF or PBS for different time points (5min to 1hour). Inhibitor compound treatment was pre-treated at 37°C for 2 hours before the first cytokine treatment, then co-treated with IL-1β and TNF for 18-24 hours. The inhibitors used in these studies are: 5uM C646 (Selleck Chemicals, #S7152), 5uM MB-3 (Sigma-Aldrich, #M2449), and 5uM MG149 (Medchem Express, #HY-15887).

### Neuronal viability assay

10,000 N2A neuronal cells (ATCC, #CCL-131) were plated in each well of 96-well plates and preactivated with 100 ng/mL mouse IFN-γ (R&D Systems, #485-MI-100) for 18-24 hours. Thereafter, medium was replaced after extensive washing with 1X PBS, adding 200 mL of astrocyte-conditioned medium per well. 18-24 hours later the supernatant was harvested for cytotoxicity evaluation by measuring LDH release with CytoTox 96 nonradioactive cytotoxicity assay kit (Promega, #G1780), followed the protocol suggested by the manufacturer.

### Lactate release measurement

2 days after seeding on 48-well plate, 200,000 astrocytes were cultured in 1mM glucose DMEM medium (#A1443001, GIBCO) supplied 100 unit/mL penicillin/streptomycin without FBS at least 48h. Thereafter, medium was removed, cells washed with 1X PBS, treated with 100 ng/mL IL-1β (R&D Systems, #401-ML-005, 100 µg/mL stock in PBS) and 50 ng/mL TNF (R&D Systems, #410-MT-010, 100 µg/mL stock in PBS) diluted in DMEM (Thermo Fisher Scientific, #11965118) that was supplemented with 10% FBS (Life Technologies, #10438026) and 1% penicillin/streptomycin (Life Technologies, #15140122) or PBS. The medium was replaced after 18-24 hours, then astrocytes were rested for 6 days. Medium was replaced every 2–3 days. At day 7, astrocytes were re-treated (2X) or treated (1X) with 100 ng/mL IL-1β and 50 ng/mL TNF or PBS for 1h. The medium was collected for filtration with Amicon ultracel 10k (Millipore, #UFC501096) under 12,000 g centrifuge at 4°C for 30 min. Lactate in flow-through medium was quantified using the L-lactate assay kit (Biomedical Research Service & Clinical Application, #A-108) and GloMax microplate reader (Promega) as suggested by the manufacturer. The absorption signal was normalized to cell number estimated using CyQUANT cell proliferation assay kit (Invitrogen, #C7026) first and then compared to control group for data analysis.

### Immunostaining of mouse CNS tissue

Mice were intracardially perfused with ice cold 1X PBS and a 0.5-cm section of the superior spinal column was excised from the mouse then post-fixed in 4% PFA overnight at 4°C. Following fixation, the vertebral column was de-calcified with 20% EDTA (pH=7.4) for one week at 4°C with inversion, then the tissue was dehydrated with 30% sucrose for one week at 4°C. Spinal cords were frozen in OCT (Sakura, #4583) and 8-20 µm sections were prepared by cryostat on SuperFrost Plus Gold slides (Fisher Scientific, #15-188-48). Sections were permeabilized with 1X permeabilization buffer (BD Biosciences, #554723) for 10 minutes at RT, then blocked using serum-free protein block (Agilent, #X0909) for 10 minutes at RT. Sections were then incubated with primary antibodies diluted in 1X permeabilization buffer overnight at 4°C. Following primary antibody incubation, sections were washed 3X with 1X permeabilization buffer and incubated with secondary or conjugated antibodies diluted in 1X permeabilization buffer for 1 hour at RT. After secondary and conjugated antibody incubation, sections were stained with 1µg/ml DAPI (Sigma-Aldrich, #D9542) diluted in 1X permeabilization buffer for 5 minutes at RT, then washed 3X with 1X permeabilization buffer and mounted with ProLong Gold Antifade Mountant (Fisher Scientific, #P36930). Iterative labeling using rabbit primary antibodies was accomplished by incubating with a single primary antibody on day 1, staining with the anti-rabbit Fab 568 fragment on Day 2, washing 6X with 1X permeabilization buffer, followed by incubation with primary and secondary antibodies as described above. Primary antibodies used in this study were: rabbit anti-p300 (Cell Signaling Technology, 1:50, #70088S), rabbit anti-ACLY (ProteinTech Group, 1:100, #15421-1-AP), rabbit anti-H3K27ac (Abcam, 1:100, #ab4729) and goat anti-SOX9 (R&D Systems, 1:100, #AF3075). Secondary antibodies used in this study were: Rhodamine Red-X-AffiniPure Fab Fragment donkey anti-rabbit IgG (H+L) (Jackson Immunoresearch, #711-297-003) and Alexa Fluor 647 AffiniPure Fab Fragment donkey anti-rabbit IgG (H+L) (Jackson Immunoresearch, #711-606-152) both at 1:1000 working dilution. The conjugated antibody used in this study was mouse anti-GFAP Alexa Fluor 488 (GA5) (Fisher Scientific, 1:100, #53-9892-82). For FluoroMyelin

Fluorescent Myelin staining, sections were permeabilized and blocked with 4% Bovine Serum Albumin (BSA, Sigma, #A3294), 0.3% Triton X-100 (Sigma, T8787) in 1X PBS for 1 hour in RT. To label the myelinated area, Fluoromyelin dye (Thermo Fisher Scientific, 1:300, #F34652) were stained in 1X permeabilization buffer (BD Biosciences, #554723) for 30 min in RT. Stained spinal cord tissues were washed 3X with 1X permeabilization buffer and mounted with ProLong Gold Antifade Mountant. Sections were imaged were stitched on an LSM-880-AiryScan confocal microscope using the ZEN Black software (Zeiss), and image analysis was performed with FIJI version of ImageJ (NIH).

### Immunostaining of CNS tissue from MS patients

For immunostaining of paraffin sections, sections were deparaffinized in 2X washes of Xylenes for 10 minutes (Sigma-Aldrich, #214736), followed by 2X washes in 100% EtOH for 10 minutes, 1X wash in 95% EtOH for 5 minutes, 1X wash in 70% EtOH for 5 minutes, 1X wash in 50% EtOH for 5 minutes, and then slides were rinsed with ddH2O for 5 minutes. Antigen retrieval was performed by placing slides in boiling Epitope Retrieval Solution (IHC World, #IW-11000) for 20 minutes. Slides were dried, a hydrophobic barrier was made (Vector Laboratories, #H-4000) and sections were washed 3X for 5 minutes with 0.3% Triton X-100 in PBS (PBS-T). Sections were blocked with 5% donkey serum (Sigma-Aldrich, #D9663) in 0.3% PBS-T at RT for 30 minutes. Sections were then incubated with primary antibody diluted in blocking buffer overnight at 4°C. Following primary antibody incubation, sections were washed 3X with 0.3% PBS-T and incubated with secondary antibody diluted in blocking buffer for 2 hours at RT. Sections were then washed 3X with 0.3%PBS-T, dried, and coverslips were mounted. Iterative labeling using rabbit primary antibodies was accomplished by incubating with a single primary antibody on day 1, staining with the anti-rabbit Fab 568 fragment on day 2, washing 6X with 1X permeabilization buffer, followed by incubation with primary and secondary antibodies as described above. Primary antibodies used in this study were: rabbit anti-P300 (Cell Signaling Technology, 1:50, #70088S), rabbit anti-ACLY (ProteinTech Group, 1:100, #15421-1-AP) and goat anti-SOX9 (R&D Systems, 1:100, #AF3075). Secondary antibodies used in this study were: Rhodamine Red-X-AffiniPure Fab Fragment donkey anti-rabbit IgG (H+L) (Jackson Immunoresearch, #711-297-003) and Alexa Fluor 647 AffiniPure Fab Fragment donkey anti-rabbit IgG (H+L) (Jackson Immunoresearch, #711-606-152) both at 1:1000 working dilution. The conjugated antibody used in this study was mouse anti-GFAP Alexa Fluor 488 (GA5) (Fisher Scientific, 1:100, #53-9892-82). Human brain tissue was obtained from patients diagnosed with clinical and neuropathological MS diagnosis according to the revised 2017 McDonald’s criteria^60^. Tissue samples were collected from healthy donors and MS patients with full ethical approval and informed consent as approved by the local ethics committee from the Centre de Recherche du Centre Hospitalier de l’Université de Montréal(CRCHUM) under ethical approval number BH07.001, Nagano 20.332-YP. Autopsy samples were preserved, and lesions classified using luxol fast blue/H&E staining as previously published^61,62^.

### Unbiased immunofluorescence quantification

Astrocyte H3K27ac, ACLY, and p300 expression were quantified as previously described^63,64^. Briefly, GFAP-positive cells were detected by implementing the *bwlabel* algorithm for labeling connected components in 2-D images using MATLAB (MathWorks). Small puncta sized < 50 μm^2^ were regarded as artifacts and were filtered out. Blood vessel related fluorescence artifacts were manually removed. Next, the averaged H3K27ac, ACLY, and p300 fluorescence intensities were calculated per individual GFAP-positive cells and outliers were removed using the Robust Regression and Outlier Removal (ROUT) method with coefficient Q = 1% ^65^.

### CRISPR/Cas9 lentivirus production

Lentiviral constructs were generated by modifying the *pLenti-U6-sgScramble-Gfap-Cas9-2A-EGFP-WPRE* lentiviral backbone, described previously^12^. This backbone contains derivatives of the previously described reagents lentiCRISPR v2 (a gift from Feng Zhang, Addgene plasmid #52961^66^), and lentiCas9-EGFP (a gift from Phil Sharp and Feng Zhang, Addgene plasmid #63592^67^). The *Gfap* promoter is the ABC_1_D *gfa2 GFAP* promoter^68^.

Substitution of sgRNAs was performed through a PCR-based cloning strategy using Phusion Flash HF 2X Master Mix (Thermo Fisher, #F548L). A three-way cloning strategy was developed to substitute sgRNAs using the following primers: U6-PCR-F 5’-AAAGGCGCGCCGAGGGCCTATTT-3’, U6-PCR-R 5’-TTTTTTGGTCTCCCGGTGTTTCGTCCTTTCCAC-3’, cr-RNA-R 5’-GTTCCCTGCAGGAAAAAAGCACCGA-3’, cr-RNA-F 5’-AAAAAAGGTCTCTACCG(N_20_)GTTTTAGAGCTAGAAATAGCAAGTT-3’, where N_20_ marks the sgRNA substitution site. The following sgRNA were designed using a combination of the Broad GPP sgRNA Designer Webtool (SpyoCas9, http://portals.broadinstitute.org/gpp/public/analysis-tools/sgrna-design), Synthego (https://design.synthego.com/#/), and cross-referenced with activity-optimized sequences contained within the Addgene library #1000000096 (a gift from David Sabatini and Eric Lander)^69^. The sgRNA sequences used are as follows, with the promoter indicated in parentheses: *sgScrmbl* - 5’-GCACTACCAGAGCTAACTCA-3’, *sgEp300 -* 5’-ACAGAATTGGGACTAACCAA-3’, *sgAcly* - 5’-GAGAGAGATTGACCCCGACG-3’, *sgAcss2* -5’-GTTTTGGGGAAACATTGCCA-3’. Amplicons were purified using the QIAquick PCR Purification Kit (Qiagen, #28104) and digested using DpnI (NEB, #R0176S), BsaI-HF (NEB, #R3535/R3733), AscI (for U6 fragment) (NEB, #R0558), or SbfI-HF (for crRNA fragment) (NEB, #R3642). pLenti backbone was cut with AscI/SbfI-HF and purified using the QIAquick PCR purification kit. Ligations into the respective backbone were performed overnight at 16°C using T4 DNA Ligase Kit (NEB, #M0202L). Ligations were transformed into NEB Stable *E. Coli* (NEB, #C3040) at 42°C and the ligation products were spread onto ampicillin selection plates. After overnight incubation at 37°C, single colonies were picked and DNA was prepared using QIAprep spin miniprep kit (Qiagen, #27104). Lentiviral plasmids were transfected into HEK293FT cells according to the ViraPower Lentiviral Packaging Mix protocol (Thermo Fisher Scientific, #K497500) and lentiviruses were packaged with pLP1, pLP2, and pseudotyped with pLP/VSVG.

Supernatant was aspirated the following day and fresh medium was added. After 2 days of incubation, lentivirus was collected and concentrated using Lenti-X Concentrator (Clontech, #631231) overnight at 4°C followed by centrifugation according to the manufacturer’s protocol. Lentiviral pellets were resuspended in 1/500 of the original volume and stored at -80°C.

### Intracranial lentivirus injection

C57Bl/6J mice at age 10-12 weeks were anesthetized using 1-3% isoflurane mixed with oxygen. Heads were shaved and cleaned using 70% ethanol and Betadine (Thermo Fisher, #19-027132) followed by a medial incision of the skin to expose the skull. The lateral ventricles were targeted bilaterally using the coordinates: +/-1.0 (lateral), -0.44 (posterior), - 2.2 (ventral) relative to Bregma. Mice were injected with approximately 10^7^ total IU of lentivirus measured by qPCR Lentivirus Titer kit (abm, #LV900) and delivered by two 10 µL injections using a 25 µL Hamilton syringe (Sigma-Aldrich, #20787) on a stereotaxic alignment system (Kopf, #1900) and the incision was sutured. Mice received 1 mg/kg Buprenorphine-SR via subcutaneous injection and were permitted to recover 7 days in a separate clean cage before induction of EAE.

### EAE induction

EAE was induced with 100 μg of MOG_35-55_ (Genemed Synthesis Inc., #110582) emulsified in freshly prepared complete Freund’s adjuvant (Incomplete Freund’s Adjuvant (BD Biosciences, #BD263910) mixed with microbacterium tuberculosis H-37Ra (BD Biosciences, #231141); final concentration 5 mg/ml). All animals received 2 subcutaneous injections of 100 μL each of MOG and a single intraperitoneal injection of 400 ng pertussis toxin (List Biological Laboratories, #180) in 200 μL of PBS. Mice received a second injection of PTx 48 hours after the initial injection. Mice were monitored and clinical scores were documented daily until the end of the experiment. Mice were sacrificed at different time points of disease. EAE clinical scores were defined as follows: 0 – no signs, 1 – fully limp tail, 2 – hindlimb weakness, 3 – hindlimb paralysis, 4 – forelimb paralysis, 5 – moribund.

### Isolation of cells from the adult CNS

Astrocytes were isolated by flow cytometry as described^10–12,14–16^ and by modifying a previously described protocol^70^. Briefly, mice were perfused with 1X PBS and the CNS was isolated into 10 mL of enzyme digestion solution consisting of 75 µL Papain suspension (Worthington, #LS003126) diluted in enzyme stock solution (ESS) and equilibrated to 37C. ESS consisted of 10 mL 10X EBSS (Sigma-Aldrich, #E7510), 2.4 mL 30% D(+)-Glucose (Sigma-Aldrich, #G8769), 5.2 mL 1M NaHCO3 (VWR, #AAJ62495-AP), 200 µL 500 mM EDTA (Thermo Fisher Scientific, #15575020), and 168.2 mL ddH2O, filter-sterilized through a 0.22 µm filter. Samples were shaken at 80rpm for 30-40 minutes at 37°C. Enzymatic digestion was stopped with 1 mL of 10X hi ovomucoid inhibitor solution and 20 µL 0.4% DNase (Worthington, #LS002007) diluted in 10 mL inhibitor stock solution (ISS). 10X hi ovomucoid inhibitor stock solution contained 300 mg BSA (Sigma-Aldrich, #A8806), 300 mg ovomucoid trypsin inhibitor (Worthington, #LS003086) diluted in 10 mL 1X PBS and filter sterilized using at 0.22 µm filter. ISS contained 50 mL 10X EBSS (Sigma-Aldrich, #E7510), 6 mL 30% D(+)-Glucose (Sigma-Aldrich, #G8769), 13 mL 1M NaHCO3 (VWR, #AAJ62495-AP) diluted in 170.4 mL ddH2O and filter-sterilized through a 0.22 µm filter. Tissue was mechanically dissociated using a 5 mL serological pipette and filtered through at 70 µm cell strainer (Fisher Scientific, #22363548) into a fresh 50 mL conical. Tissue was centrifuged at 500g for 5 minutes and resuspended in 10 mL of 30% Percoll solution (9 mL Percoll (GE Healthcare Biosciences, #17-5445-01), 3 mL 10X PBS, 18 mL ddH2O). Percoll suspension was centrifuged at 500g for 25 minutes with no brakes. Supernatant was discarded and the cell pellet was washed 1X with 1X PBS, centrifuged at 500g for 5 minutes and prepared for downstream applications.

### Isolation of mouse splenic cells

Spleens were isolated prior to mouse perfusion and mechanically dissociated. Red blood cells were lysed with ACK lysing buffer (Life Technology, #A10492-01) for 2 min and washed with 0.5% BSA, 2 mM EDTA pH=8.0 in 1X PBS and prepared for downstream applications.

### FACS

Cells were stained in the dark on ice for 20 minutes with FACS antibodies. Cells were then washed once with 1X PBS and resuspended in 1X PBS for sorting as described^10–12,14–16^. Antibodies used in this study were: PE anti-mouse CD45R/B220 (BD Biosciences, #553089, 1:100), PE anti-mouse TER-119 (Biolegend, #116207, 1:100), PE anti-O4 (R&D Systems, #FAB1326P, 1:100), PE anti-CD105 (eBioscience, #12-1051-82, 1:100), PE anti-CD140a (eBioscience, #12-1401-81, 1:100), PE anti-Ly-6G (Biolegend, #127608, 1:100), BV786-Ly6C (BD Biosciences, #740850, 1:100), APC anti-CD45 (eBioscience, #17-0451-83, 1:100), APC-Cy7 anti-CD11c (BD Biosciences, #561241, 1:100), and FITC anti-CD11b (eBioscience, #11-0112-85, 1:100). All cells were gated on the following parameters: CD105^neg^CD140a^neg^O4^neg^Ter119^neg^Ly-6G^neg^CD45R^neg^. Astrocytes were gated on: CD11b^neg^CD45^neg^Ly-6C^neg^CD11c^neg^. Microglia were subsequently gated on: CD11b^high^CD45^low^Ly-6C^low^. Pro-inflammatory monocytes were gated on: CD11b^high^CD45^high^Ly-6C^high^. Compensation was performed on single-stained samples of cells and an unstained control. Cells were sorted on a FACS Aria IIu (BD Biosciences) and analyzed by BD FACSDIVA (v.8.0.1).

### FACS analysis of T cells

CNS and splenic cell suspensions were stimulated with 50 ng/mL phorbol 12-myristate 13-acetate (PMA, Sigma-Aldrich, #P8139), 1 µM Ionomycin (Sigma-Aldrich, # I3909-1ML), GolgiStop (BD Biosciences, #554724, 1:1500) and GolgiPlug (BD Biosciences, #555029, 1:1500) diluted in T cell culture medium (RPMI (Life Technologies, #11875119) containing 10% FBS, 1% penicillin/streptomycin, 50 µM 2-metcaptoethanol (Sigma-Aldrich, #M6250), and 1% non-essential amino acids (Life Technologies, #11140050). After 4 hours, cell suspensions were washed with 0.5% BSA, 2 mM EDTA in 1X PBS and incubated with surface antibodies and a live/dead cell marker on ice. After 30 min, cells were washed with 0.5% BSA, 2mM EDTA in 1X PBS and fixed according to the manufacturer’s protocol of an intracellular labeling kit (eBiosciences, #00-5523-00). Surface antibodies used in this study were: BUV661 anti-mouse CD45 (BD Biosciences, #565079, 1:100), BV750 anti-mouse CD3 (Biolegend, #100249, 1:50), PE-Cy7 anti-mouse CD4 (eBioscience, #25-0041-82, 1:100), BV785 anti-mouse CD11b (BD Biosciences, #101243, 1:100), BV570 anti-mouse Ly6C (Biolegend, #128030, 1:100), BUV805 anti-mouse CD8a (BD Bioscience, #564920, 1:100), BUV563 anti-mouse Ly6G (BD Biosciences, #565707, 1:100), BUV737 anti-mouse CD11c (BD Biosciences, #612797, 1:100), Pe/Cy5 anti-mouse CD44 (BioLegend, #103010, 1:100), Intracellular antibodies were: APC anti-mouse IFN-γ (BD Biosciences, #554413, 1:100), PE anti-mouse IL-17A (eBiosciences, #12-7177-81, 1:100), BV421 anti-mouse GM-CSF (BD Biosciences, #564747, 1:100), FITC anti-mouse FoxP3 (eBiosciences, #11-5773-82, 1:100). Gating of CNS and splenic cells was performed on >10,000 live CD4^+^ cells. FACs was performed on a Symphony A5 (BD Biosciences).

### Isolation, culture, and stimulation of human primary astrocytes

Primary human fetal astrocytes were isolated as previously described^71–73^. Second trimester fetal brain tissues were obtained from Centre Hospitalier Universitaire Sainte-Justine, Montreal, Canada or from the Birth Defects Research Laboratory, University of Washington, Seattle, USA with maternal written consent and under local ethic boards’ approval. Isolation of glial cells was carried out through mechanical and chemical digestion. Primary fetal astrocytes were maintained in Dulbecco’s Modified Eagle Medium (DMEM)/F12 (Sigma-Aldrich, #D8437) with 10% fetal bovine serum (FBS, Wisent Bio Products, #080-150), 1% penicillin/streptomycin (ThermoFisher, #15140122) and 1% Glutamax (ThermoFisher, #35050-061) and grown in incubators set to 37°C with 5% CO2. For in vitro conditions, human primary astrocytes were seeded at 50,000 cells per well. The cell media was aspirated, cells were washed with 37°C PBS and the fresh, appropriate media was then added into the wells. The cells were treated for 18-24 hours with 10 ng/mL IL-1β (Gibco, #PHC0815) and 5 ng/mL TNF (Gibco, #PHC3016) in basal media. After the first stimulation, cells were washed 2 times with warmed PBS and then cultured in normal culture media for the subsequent 6 days. At day 7, human primary astrocytes were re-treated (2X) or treated (1X) with 10 ng/mL IL-1β and 5ng/mL TNF or PBS for 2 hours. For the control conditions, the media aspiration, media change and PBS washing occurred alongside all treated cells. At the end of these treatments, total RNA was isolated, transcribed and subjected to qPCR. Inhibitor compound treatment was pre-treated at 37°C for 2 hours before the first cytokine treatment, then co-treated with IL-1β and TNF for 18-24 hours. The inhibitor used in these studies is: 5uM C646 (Selleck Chemicals #S7152).

### RNA expression analysis of primary astrocytes

Primary astrocytes were lysed in Buffer RLT (Qiagen) and RNA was isolated from cultured astrocytes using the Qiagen RNeasy Mini kit (Qiagen, #74106). cDNA was transcribed using the High-Capacity cDNA Reverse Transcription Kit (Life Technologies, #4368813). Gene expression was then measured by qPCR using Taqman Fast Universal PCR Master Mix (Life Technologies, #4367846). Taqman probes used in this study are: *Actb* (Mm02619580_g1), *Nos2* (Mm00440502_m1), *Ccl2* (Mm00441242_m1), *Il6* (Mm00446190_m1), *Ep300* (Mm00625535_m1), *Kat5* (Mm01231512_m1), *Kat2b* (Mm00517402_m1), *Acly* (Mm01302282_m1), *Acss2* (Mm00480101_m1), *ACTB* (Hs01060665_g1), *NOS2* (Hs01075529_m1), *IL6* (Hs00174131_m1), and *CCL2* (Hs00234140_m1). qPCR data were analyzed by the ddCt method by normalizing the expression of each gene for each replicate to β-actin and then to the control group.

### Chromatin immunoprecipitation (ChIP)

Approximately >1 million astrocytes were used by cell preparation according to the ChIP-IT Express Enzymatic Shearing and ChIP protocol (Active Motif, #53009). Briefly, cells were fixed in 1% formaldehyde for 10 minutes with gentle agitation, washed in 1X PBS, washed for 5 minutes in 1X glycine Stop-Fix solution in PBS, and scraped in 1X PBS supplemented with 500 µM PMSF. Cells were pelleted, nuclei isolated, and chromatin sheared using the Enzymatic Shearing Cocktail (Active Motif) for 10 minutes at 37°C with vortexing every 2 minutes. Sheared chromatin was immunoprecipitated according to the Active Motif protocol overnight at 4°C with rotation. The next day, the protein-bound magnetic beads were washed 1X with ChIP buffer 1, 1X with ChIP buffer 2, and 1X with 1X TE. Cross-links were reversed in 100 µL of 0.1% SDS and 300 mM NaCl in 1X TE at 63°C for 4-5 hours, as described ^74,75^. DNA was purified using Agencourt AMPure XP Beads (Beckman Coulter, #A63881). qPCR was performed using Fast SYBR Green Master Mix (Thermo Fisher Scientific, #4385612). Anti-IgG immunoprecipitation and input were used as controls. We used rabbit anti-p300 (Genetex, GTX114541, 1:100), rabbit anti-H3K27ac (Abcam, 1:100, #ab4729), and rabbit IgG polyclonal isotype control (Cell Signaling, 2975S, 1:100). PCR primers were designed with Primer3^76^ to generate 50-150 bp amplicons. Primer sequences used: p300-Il6-F: 5’-GCTGTTTCAGCTGCCTTTTT-3’, p300-Il6-R: 5’-AAAGCCGGTTGATTCTTGTG-3’, p300-Ccl2-F: 5’-GGCCCGTTAGCTGTCTGTTA-3’, p300-Ccl2-R: 5’-AGTGGTTGGAACCCTTCACA-3’, p300-Nos2-F: 5’-AACACGAGGCTGAGCTGACT-3’, p300-Nos2-R: 5’-CACACATGGCATGGAATTTT-3’, p65-Il6-F: 5’-AAGCACACTTTCCCCTTCCT-3’, p65-Il6-R: 5’-GCTCCAGAGCAGAATGAGCTA-3’, p65-Ccl2-F: 5’-CCAAATTCCAACCCACAGTT-3’, p65-Ccl2-R: 5’-AGTGAGAGTTGGCTGGTGCT-3’, Data were analyzed by ddCt relative to Input control.

### Bulk RNA-sequencing

Sorted cells were lysed and RNA isolated using the Qiagen RNeasy Micro kit (Qiagen, #74004) with on-column DNase I digestion (Qiagen, #79254). RNA was suspended in 10 µL of nuclease free water at 0.5-1 ng/µL and sequenced using SMARTSeq2^77^ at the Broad Institute Technology Labs and the Broad Genomics Platform. Processed RNA-seq data was filtered by removing genes with low read counts. Read counts were normalized using TMM normalization and CPM (counts per million) were calculated to create a matrix of normalized expression values. The fastq files of each RNA-seq data sample were aligned to the *Mus musculus* GRCm38 transcriptome using Kallisto (v0.46.1), and the same software was used to quantify the alignment results. Differential expression analysis was conducted using DESeq2, and the log2 fold change was adjusted using apeGLM for downstream analysis.

### ChIP-seq

Approximately 500,000 cells were fixed, nuclei collected, and chromatin isolated using the ChIP-IT Express Enzymatic Shearing Kit (Active Motif, #53009). Chromatin complexes were immunoprecipitated using a rabbit anti-Histone H3 (acetyl K27) antibody (Abcam #ab4729, 1:100) and rabbit IgG polyclonal isotype control (Cell Signaling, 2975S, 1:100). Cross-links were reversed in 100 µL of 0.1% SDS and 300 mM NaCl in 1X TE at 63°C for 4-5 hours, as described ^74,75^. DNA was purified using Agencourt AMPure XP Beads (Beckman Coulter, #A63881). DNA was analyzed using a 2100 Bioanalyzer (Agilent) and High Sensitivity DNA Kit (Agilent Technologies, #5067-4626). DNA libraries were then prepared for sequencing using the NEBNext Ultra II DNA Library Prep Kit (New England Biolabs, #E7645S) and sequencing adaptors were ligated using NEBNext Multiplex Oligos for Illumina (#E7500S, #E7335S) as described. ChIPed DNA was amplified by using 14 cycles according to the NEBNext protocol. DNA libraries were not size selected, but primer dimers were removed via purification using Agencourt AMPure XP Beads (Beckman Coulter, #A63881). DNA libraries consisting of ChIPed DNA and IgG control DNA for each sample were sequenced on an Illumina HiSeq 4000 using 2×75 paired end sequencing.

Raw fastq files were first checked for quality using Multiqc sequence analysis. Reads were aligned to the mouse genome (mm10) using bowtie2 (2.3.4.3)^78^. Subsequently, SAM files were sorted and filtered using Sambamba (v.0.8.2)^79^. We discarded unmapped and duplicate fragments. Sorted BAM files were indexed using samtools (v.1.15.1)^80^. BAM files were then converted to BigWig files for visualization using deeptools (v.3.5.0)^81^. Normalization for all BigWig files were carried out against effective genome size (2652783500 for mice H3K27ac ChIP-seq). Peak analysis was carried out with Macs2 (v.2.2.7)^82^ using stringent parameters (-f BAMPE --mfold 5 50 -p 0.001). A master-peak bed file was created using bedtools merge function. Significant peaks present in one sample were considered. Aligned and filtered counts were extracted from BAM files using master-peak bed file and differential peaks between p300 and scramble were determined using two-sided t-test with *p*-value < 0.05 **(Supp. Table 5)**. Specific peaks were visualized using IGV (v.2.11.4), exported as .png, and edited in Adobe Illustrator. Peak plots were visualized using deepTools computeMatrix with 500bp from peak center.

### ATAC-seq

Sequencing libraries were prepared largely as described previously^74,83,84^. After isolation of nuclei, transposition was performed using the kit (Illumina, #FC-121-1030). DNA was then amplified using NEBNext High Fidelity 2X PCR Master Mix (New England Biolabs, #M0541S) for 5 cycles. DNA quantity was then measured using a Viia 7 Real-Time PCR System (Thermo Fisher Scientific) and the number of cycles required to achieve 1/3 of maximal SYBR Green fluorescence was determined and libraries were amplified accordingly. TruSeq adaptors (universal: Ad1_noMX and barcoded: Ad2.1-Ad2.24) were used according to the Buenrostro protocol. Libraries were purified using MiniElute PCR Purification Kit (Qiagen, #28006) followed by double sided Agencourt AMPure XP bead purification (Beckman Coulter, #A63881) to remove primer dimers and large DNA fragments. Libraries were analyzed on a 2100 Bioanalyzer (Agilent Technologies) and High Sensitivity DNA Kit (Agilent Technologies, #5067-4626). Libraries were sequenced by Genewiz on an Illumina HiSeq 4000 by 2×75 bp paired end sequencing.

Paired-end ATAC-Seq reads were first assessed using FASTQC. Reads were trimmed using cutadapt to remove adapters and low quality reads below 30. Pair-end reads were then aligned against the GRCm39/mm10 mouse genome assembly with Bowtie (v2.3.0)84 in local mode, sensitive settings, and a maximum fragment size of 2000. Duplicated reads were marked using Picard (v.2.5.0). Alignments were filtered with SAMtools (v1.3) to exclude reads with mapping quality <30, not properly paired, duplicated, aligned to mitochondrial genome, and/or aligned to ENCODE blacklist regions. Alignments with an insertion size of >100bp were removed to enrich for nucleosome-free reads. ATAC-Seq peaks were called for each replicate using MACS2, using -- format BAMPE and --keep-dup all. IDR (v2.0.2) was used to determine consistency of peak detection between individual replicates and peaks with a threshold below 0.10 were merged between replicates for downstream analysis. For differential peaks, merged peaks were mapped to specific genic regions using bedtools intersect and reads were counted using subread featureCounts (v.1.6.2) to produce a count matrix. DESeq2 was then used to find differential peaks.

### FIND-seq

The transcriptome of *Acly*^+^*Ep300*^+^ astrocyte subsets was isolated and sequenced using the FIND-seq/Smart-seq3 protocol^29,85^. The process started with the preparation of oligo dT primer-conjugated agarose. 50 µM of acrydite-modified primers (/5Acryd/TTTTTTACGAGCATCAGCAGCATACGAT30V, IDT) were mixed with a 0.5% (wt/vol) allyl agarose solution (SFR allyl agarose, Lucidant Polymers), and 0.1% (wt/vol) APS (Promega, #V3131) and 0.1% (vol/vol) TEMED (Invitrogen, #15524-010) were introduced at the reaction initiation and after a 4 hour incubation at 45°C to conjugate Oligo dT primer on the agarose chain. For optimal conjugation, the reaction was continued for an additional 16 hours. Extra ultra-low gelling temperature agarose type IX-A (Sigma-Aldrich, #A2576) was used to raise total agarose concentration to 2%. The conjugated agarose gel was washed in nuclease-free water to remove unreacted primers and residual chemicals. The concentration of the oligo dT primer on the agarose was quantified using the Qubit ssDNA assay kit (Invitrogen, #Q10212) and normalized to 8 µM.

Astrocytes were resuspended at a density of 6.11 × 10^6^/mL in a buffer composed of 1% (vol/vol) Pluronic F-68 (Thermo Fisher, #24040032) and 18% (vol/vol) Opti-prep (Sigma-Aldrich, #D1556-250ML) in HBSS buffer without calcium or magnesium (Gibco, #14170112), then filtered through a 70-µm cell strainer (Fisher Scientific, #22363548) and loaded into a 1mL syringe (BD Biosciences, #309628). A lysis buffer, containing 20 mM Tris-HCl (pH 7.5) (Teknova, #T5075), 1000 mM LiCl (Sigma-Aldrich, #L7026), 1% (wt/vol) LiDS (Sigma-Aldrich, #L4632-25G), 10 mM EDTA (VWR, #E177), 10 mM DTT (Sigma-Aldrich, #43816), and 2 µg/µL of Proteinase K (New England Biolabs, #P8107S), was prepared and loaded into a 3 mL syringe (BD Biosciences, #309657). Primer-conjugated agarose was completely melted at 95°C for over 2 hours and subsequently loaded into a 3 mL syringe, which was placed into a custom-built syringe heater to maintain its molten state at 90°C. Single cells, alongside the primer-conjugated agarose and lysis buffer, were encapsulated using a Bubble-triggered device **(Extended Data Fig. 8b)**. The cell solution, lysis buffer, molten agarose solution, Droplet Generation Oil for Probes (Bio-rad Laboratories, #1864110), and pressurized air lines were connected to the Bubble-triggered device via PE/2 tubing (Scientific Commodities, #BB31695-PE/2). The flow rates were adjusted to produce 55-µm droplets: Cell: 600µL/hr, Lysis buffer: 600μL/hr, Molten primer-conjugated agarose: 1200μL/hr, Droplet Generation Oil for Probes: 5000µL/hr, Pressured air: 20psi. Following encapsulation, the droplets were incubated at 55°C for 2 hours and subsequently cooled at 4°C for 1 hour to solidify the agarose within the droplets. The hardened agarose beads were collected through a droplet-breaking procedure. Residual oil at the bottom of the tube was discarded, and the emulsion was effectively disrupted using 20% (vol/vol) 1H,1H,2H,2H-Perfluoro-1-octanol (PFO) (Sigma-Aldrich, #370533-25G) in HFE-7500 (3M™ Novec™ 7500 Engineered Fluid). The mixture was centrifuged for 5 minutes at 2,000 g at 4°C, and the OFP phase at the bottom was removed. The agarose beads were then resuspended in a buffer containing 20 mM Tris-HCl (pH 7.5), 500 mM LiCl, 0.1% (wt/vol) LiDS, and 0.1 mM EDTA. Following another centrifugation for 5 minutes at 2,000g at 4°C, the supernatant was discarded. The beads were subsequently resuspended in a buffer composed of 20 mM Tris-HCl (pH 7.5) and 500 mM NaCl (Thermo Fisher Scientific, # AM9759).

Released mRNA from the cells was captured through hybridization with the conjugated oligo dT primer. Subsequent reverse transcription (RT) leads to the formation of cDNA on the agarose bead. For the RT reaction, agarose beads were combined with a master mix comprising 1 mM dNTPs (Thermo Scientific, #R1121), 2 µM Smart-seq3 TSO (AAGCAGTGGTATCAACGCAGAGTACATrGrGrG, IDT), 6 mM MgCl_2_ (Sigma-Aldrich, #M1028), 1 M betaine (Sigma-Aldrich, #61962), 7.5% PEG-8000 (Promega, #V3011), 2 U/µL Maxima H-minus reverse transcriptase, and 0.5 U/µl RNAse inhibitor (Lucigen, #30281). The RT reaction was executed by rotating for 30 min at room temperature, followed by further incubation for 1h 30 min at 42°C. After RT, the agarose beads were washed five times in a centrifuge with a solution of 0.1% Tween-20 (Sigma-Aldrich, #P9416).

The cDNA on the agarose bead was validated via whole transcriptome amplification (WTA). The genomic material trapped in the agarose bead was stained with SYBR green I (Sigma, #S9430), and the concentration of genome-captured beads was measured using a hemocytometer, which counted the SYBR green I-stained agarose beads **(Extended Data Fig. 8e)**. Based on the calculated concentration, 750 genome-capture beads were resuspended in 25 µL of WTA master mix; this contained 1× KAPA HiFi master mix (Roche, #KK2601), 0.5 μM Smart-seq3 forward primer (TCGTCGGCAGCGTCAGATGTGTATAAGAGACAGATTGCGCAATG, IDT), and 0.1 µM Smart-seq3 reverse primer (ACGAGCATCAGCAGCATACG, IDT). The reaction underwent thermocycling: 95°C for 3 min, followed by 18 cycles of (98°C for 15s, 67°C for 20s, 68°C for 4min), then 72°C for 5min, and held at 4°C. The WTA product was extracted using 0.8× volume of AMPure XP beads (Beckman Coulter, #A63881) following the manufacturer’s protocol. DNA was measured with the Qubit dsDNA HS Assay kit. The size distribution of 1 ng of WTA product, based on the Qubit results, was confirmed on a Bioanalyzer **(Extended Data Fig. 8f)**.

The genome and transcriptome-capturing agarose beads were reintroduced into droplets to execute multiplexed droplet digital PCR (ddPCR). The PCR master mix (TaqPath ProAmp Master Mix; Thermo Fisher Scientific, #A30866) was combined with taqman probes for *Ep300* (Mm00625535_m1) and *Acly* (Mm01302282_m1), alongside 0.1% Tween-20 and 2.5% PEG 8,000 to ensure droplet stability. This bead reinjection was made by a microfluidic device, the Re-injector device **(Extended Data Fig. 8c)**. The agarose beads were soaked in the PCR master mix for 30 minutes at room temperature, centrifuged for 3 minutes at 2000g at 4°C, and the supernatant was discarded. The bead pellet was loaded into a 1 mL syringe. A PCR master mix without beads was loaded into a 1mL syringe, and Droplet Generation Oil for EvaGreen (Bio-rad Laboratories, #1864112) was loaded into a 10mL syringe. The bead, PCR master mix, and oil were connected to the Re-injector device via PE/2 tubing. The flowrates were optimized to generate a single bead per 80-µm sized droplet with PCR master mix; Bead: 200μL/hr, PCR master mix: 300μL/hr, Oil: 1800 μL/hr. Droplets containing the reinjected bead were collected in a PCR tube and underwent thermocycling to generate TaqMan signals. The thermocycling conditions were set as follows: 88°C for 10 min, followed by 55 cycles of (88°C for 30 s, 59°C for 1 min), then 72°C, and held at 59°C.

After thermocycling, the emulsion was loaded into a 3 mL syringe. The detection and sorting of droplets were performed using an in-house droplet sorting system^85^, and a droplet sorter device **(Extended Data Fig. 8d)**. Two 30 mL syringes filled with HFE-7500 oil and a 3 mL syringe with Droplet Generation Oil for EvaGreen were prepared. A 2M NaCl solution filled the electrode and moat channels using a 10mL syringe, and a high voltage amplifier was connected to the needle of the syringe linked to the electrode channel. The droplet sorter device was connected to the emulsion, drop spacing oil (Droplet Generation Oil for EvaGreen in 3mL syringe), additional spacer oil (HFE-7500 in 30 mL syringe), bias oil (HFE-7500 in syringe) and pressurized air via PE/2 tubing. The assigned flowrates were as follows: Emulsion: 100μL/hr, Droplet spacing oil: 300μL/hr, additional spacing oil Oil: 3000μL/hr, Bias oil: 4000μL/hr, Pressured Air: 20psi. The TaqMan signals were excited using two lasers (OptoEngine, #MLL-FN-473, #MLL-III-532), and detected with bandpass filters (GFP channel FF552 and FF01-517/20; RFP channel FF593 and FF01-572/28 and photomultiplier tubes (PMTs) (Thorlabs, #PMM01, #PMM02) setup. Negative, singles, and double positive droplets were sorted directly into a 1.5mL tube in aliquot of 100 drops, and 50µL of nuclease-free water was added on the oil phase.

The 1.5 mL tube containing sorted droplets was incubated at -80°C for 2 hours, then thawed at 60°C for 10 minutes to facilitate droplet breaking and release of the cDNA conjugated to the agarose. The oil phase at the bottom was discarded, and the cDNA-containing water phase was harvested and transferred to a PCR tube. For the amplification of cDNA, a PCR master mix was added to the PCR tube. This mix contained 0.3 mM dNTP, 1× KAFA HiFi enzyme and buffer (Roche, #KK2502), 0.5 µM Smart-seq3 forward primer and 0.1 µM Smart-seq3 reverse primer. The reaction was subjected to the following thermocycling conditions: 95°C for 3 min, followed by 24 cycles of (98°C for 15 s, 67°C for 20 s, 68°C for 4 min), then 72°C for 5 min, and held at 4°C. The amplified material was cleaned up with 0.8× volume of AMPure XP beads and eluted in 10 µL of nuclease-free water. Finally, the DNA library was prepared using the Nextera XT Library Prep Kit (Illumina, #FC-131-1024), following the manufacturer’s protocol.

### Pathway analysis

GSEA or GSEAPreranked analyses were used to generate enrichment plots for bulk RNA-seq or scRNA-seq data^86,87^ using MSigDB molecular signatures for canonical pathways: KEGG/Reactome/Biocarta (c2.cp.all), Motif (c3.all), Gene ontology (c5.cp.all), and Hallmark (h.all). To determine regulators of gene expression networks, Ingenuity Pathway Analysis software (Qiagen) was used by inputting gene expression datasets with corresponding log (FoldChange) expression levels compared to other groups. “Canonical pathways” and “upstream analysis” metrics were considered significant at *p*-value <0.05. Gene sets used for signature scores are indicated in the corresponding figure. The RNA-Seq samples were processed using the following pipeline: FASTQC was used to assess the read quality, and then Kallisto was used to do the pseudo-alignment and quantification. DESeq2 was used to find the differentially expressed genes. From the differential expression analysis, the log2FoldChange of the genes were multiplied by the -log10 (adjusted *p*-value), and the products were ordered in a decreasing order, which formed an elbow plot and knee plot. Genes that are before the elbow point and after the knee points formed the astrocyte epigenetic memory signatures (up and down). Independent component analysis (ICA) was used to analyze the FIND-Seq dataset to extract the combined signatures. Previously published datasets were analyzed from the following studies at their respective accession numbers^10,30,31^. Seurat (v3) was used to analyze the single cell RNA-Seq dataset. The function AddModuleScore() is used to quantify the scores of signatures derived from bulk RNA-Seq analyses for every cell. Monocle (v2) was then used to construct the pseudo-time trajectory for the single cell RNA-Seq dataset.

## Data and materials availability

Sequencing data were deposited into GEO under the SuperSeries accession number GSE237558. The Reviewer access token is: *Yfcfyqwwnfetpud*. All other materials will be made available upon request.

## Acknowledgments

This work was supported by grants NS102807, ES02530, ES029136, AI126880 from the NIH; RG4111A1 and JF2161-A-5 from the NMSS; RSG-14-198-01-LIB from the American Cancer Society; and PA-1604-08459 from the International Progressive MS Alliance. H.-G.L. was supported by a Basic Science Research Program through the National Research Foundation of Korea (NRF) funded by the Ministry of Education (2021R1A6A3A14039088). J-H Lee was supported by Basic Science Research Program funded by National Research Foundation of Korea(NRF)/Ministry of Education (2022R1A6A3A03071157), and long-term postdoctoral fellowship funded by Human Frontier Science Program (LT0015/2023-L). T.I. was supported by the EMBO postdoctoral fellowship (ALTF: 1009-2021). G.P. is a trainee in the Medical Scientist Training Program funded by NIH T32 GM007356. The content of this manuscript is solely the responsibility of the authors and does not necessarily represent the official views of the National Institute of General Medical Science or NIH. M.A.W. was supported by NINDS, NIMH, and NCI (R01MH132632, R01MH130458, R00NS114111, T32CA207201). A.P. holds the T1 Canada Research Chair in MS and is funded by the Canada Institute of Health Research, the NMSS, and the Canadian Foundation for Innovation. I.C.C. was supported by K22AI152644 and DP2AI154435 from the NIH. We thank L. Li for technical assistance and all members of the Quintana laboratory for helpful advice and discussions. We thank R. Krishnan for technical assistance with flow cytometry. We thank BDRL members Kimberly A. Aldinger, Dan Doherty, Ian G. Phelps, Jennifer C. Dempsey, Kevin J. Lee, and Lucinda A. Cort.

## Authors’ contribution

H-G.L., V.R., M.A.W., and F.J.Q. designed research; H-G.L., J.M.R., C.F.A., J-H.L., L.E.F., F.P., C-C.C., L.S., T.I., F.G., M.C., L.M.S., J.E.K., G.P., S.E.J.Z., and V.R. performed experiments; H-G.L., Z.L., C.F.A., K.K., and F.J.Q. performed bioinformatic analyses; S.W.S. performed FIND-seq; J.A., A.P., and I.C.C. provided unique reagents and discussed and/or interpreted findings; H-G.L., and F.J.Q. wrote the paper with input from coauthors; F.J.Q. directed and supervised the study.

